# Nonallosteric Sirtuin Enzyme Activation: Characterization of Hit Compounds

**DOI:** 10.1101/2020.04.17.045187

**Authors:** A Upadhyay, X Guan, S Munshi, R Chakrabarti

## Abstract

Mammalian sirtuins (SIRT1-SIRT7) are a family of nicotinamide adenine dinucleotide (NAD^+^)-dependent protein deacylases that play critical roles in lifespan and age-related diseases. The physiological importance of sirtuins has stimulated intense interest in designing sirtuin-activating compounds. However, except for allosteric activators of SIRT1-catalyzed reactions that are limited to particular substrates, a general framework for the design of sirtuin-activating compounds has been lacking. Recently, we introduced a general mode of sirtuin activation that is distinct from the known modes of enzyme activation, establishing biophysical properties of small molecule modulators that can, in principle, result in enzyme activation for various sirtuins and substrates. Here, we characterize small molecules reported in the literature to activate the SIRT3 enzyme using a variety of computational, biochemical and biophysical techniques including protein-ligand docking, molecular dynamics simulation, nonlinear reaction dynamics simulation, kinetic assays and thermodynamic assays with multiple substrates and protocols. In particular, we identify the mechanism of action of the compound honokiol on the human SIRT3 enzyme, modeling its effect on active site conformational degrees of freedom and demonstrating how it nonallosterically activates the human SIRT3 enzyme under physiologically relevant conditions. We show that honokiol constitutes a hit compound for the design of a new generation of nonallosteric activators that can activate SIRT3 through the proposed mechanism-based mode of activation.

## INTRODUCTION

Sirtuin (silent information regulator) enzymes, which catalyze NAD^+^-dependent protein post-translational modifications, have emerged as critical regulators of many cellular pathways including age-related diseases [1]. Sirtuins are NAD^+^-dependent lysine deacylases, requiring the cofactor NAD^+^ to cleave acyl groups from lysine side chains of their substrate proteins, and producing nicotinamide (NAM) as a by-product. The mechanism of sirtuin-catalyzed deacylation is depicted in Fig. 1 [2-7].

**Fig. 1.**
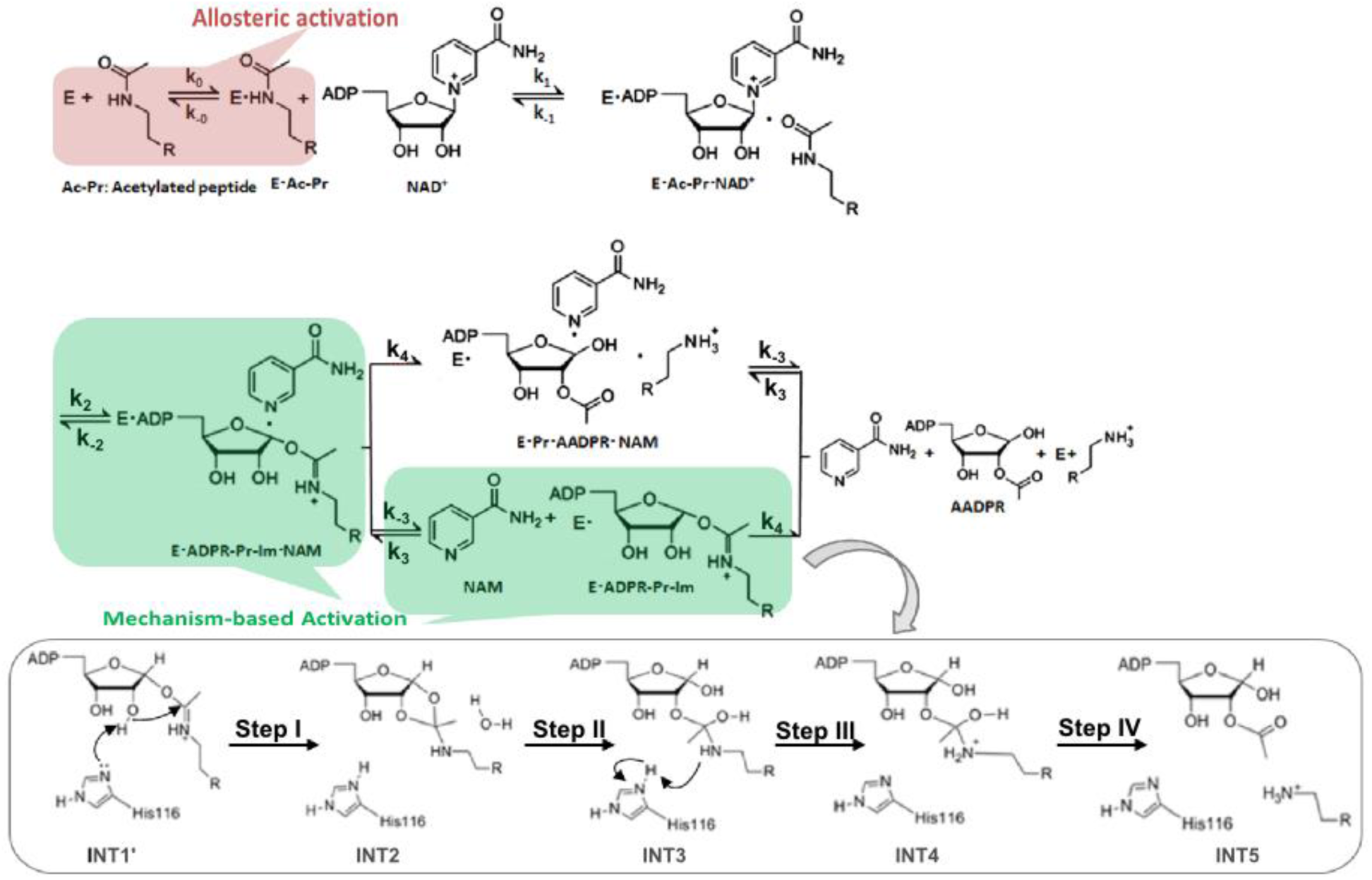
Chemical mechanism of sirtuin-catalyzed deacylation and modes of sirtuin activation. Following sequential binding of acylated peptide substrate and NAD^+^ cofactor, the reaction proceeds in two consecutive stages: i) cleavage of the nicotinamide moiety of NAD^+^ (ADP-ribosyl transfer) through the nucleophilic attack of the acetyl-Lys side chain of the protein substrate to form a positively charged O-alkylimidate intermediate, and ii) subsequent formation of deacylated peptide. For simplicity, all steps of stage ii as well as AADPR + Pr dissociation are depicted to occur together with rate limiting constant *k*_*4*_. The schematic highlights mechanism-based activation through NAD^+^ *K*_*m*_ reduction rather than the *K*_*d*_ peptide reduction that known allosteric sirtuin activators elicit.

The search for pharmacological agents that activate sirtuins has become a focus for many anti-aging studies due to sirtuins’ lifespan extending effects in mammals [8, 9]. Almost all reported sirtuin activators target SIRT1 through allosteric activation [9-13]. These allosteric activators reduce the dissociation constant of the substrate (e.g. the acylated protein dissociation constant *K*_*d, Ac*−Pr_ in the case of sirtuins) to activate the enzyme. Allosteric activators of SIRT1 bind outside of the active site to an allosteric domain that is not shared by SIRT2-7 [11]. Moreover, allosteric activators only work with a limited set of SIRT1 substrates [14-18].

Allosteric activation is one of four known modes by which small molecules can activate enzymes [10]. Aside from allosteric activation, enhancement of enzymatic activity via “derepression of inhibition” has been explored theoretically and experimentally. Theoretical models proposed have been limited to inhibitors that are exogenous to the reaction, and experimental studies have considered alleviation of product inhibition through competition with product binding [2]. These approaches can only enhance enzyme activity in the presence of inhibitor or product accumulation and hence are not included among the four known modes of enzyme activation.

Small molecules that can activate sirtuins through modes of action other than the reduction of substrate *K*_*d*_ are of particular interest, because sirtuin activity often declines during aging for reasons other than substrate depletion. Unlike allosteric activators like resveratrol, which are SIRT1-specific and have not been successfully applied to other sirtuins [11], NAD^+^ supplementation [19] can activate most mammalian sirtuins in a substrate-independent fashion. On the other hand, the effects of NAD^+^ supplementation are not specific to sirtuins and prohibitively high concentrations of NAD^+^, along with associated undesirable side effects, may be required to elicit the increases in sirtuin activity required to combat age-related diseases [20].

A preferred general strategy for activation of sirtuins (Fig. 1) would be to lower the *K*_*m*_ for NAD^+^ (*K*_*m, NAD*+_). *K*_*m, NAD*+_ reduction would have a similar activating effect to NAD^+^ supplementation, but would be selective for sirtuins and could potentially even provide isoform-specific sirtuin activation. Unlike allosteric activation that reduces *K*_*d, Ac*−Pr_, this approach could be applicable to multiple sirtuins and substrates. Moreover, since NAD^+^ is directly involved in the sirtuin ADP ribosylation reaction, regulation of *K*_*m, NAD+*_ may theoretically be achievable by methods including but not limited to varying the binding affinity of NAD^+^.

Several compounds have been reported in the literature as being activators of sirtuins other than SIRT1 [21-24]. Concurrently, sirtuins whose preferred substrates are long chain acylated peptides were reported to have their activities on shorter chain acyl groups enhanced in the presence of certain fatty acids, further corroborating the feasibility of substrate-specific sirtuin activation [25, 26]. However, some of the above compounds have been studied using only endpoint assays [21, 23] and others have been evaluated using only labeled assays [23] that may lead to false positives. Moreover, studies that have employed label-free initial rate assays [24] have not shed light on the underlying mechanism by which activation may occur.

In recent work [27], we presented a general framework for activation of sirtuin enzymes that is distinct from any of the known modes of enzyme activation. We introduced a steady-state model of sirtuin-catalyzed deacylation reactions in the presence of NAD^+^ cofactor and endogenous inhibitor NAM, and then established quantitatively how k_cat_*/K*_*m,NAD+*_ can be modified by small molecules, identifying the biophysical properties that small molecules must have to function as such mechanism-based activators. In this paper, we apply this theory of mechanism-based enzyme activation to experimentally and computationally characterize several small molecules that have been reported to be activators of the SIRT3 enzyme in order to determine whether these compounds possess the proposed characteristics of mechanism-based sirtuin activating compounds (MB-STACs). We also extend this framework to include the modulation of active site structure by mechanism-based activators, as well as activation under physiologically relevant non-steady state conditions. This constitutes the first application of the theory for mechanism-based enzyme activation to experimental and computational data, and as such provides a foundation for drug development of MB-STACs. We employ our approach to characterize the effects of the compound honokiol, which was reported to be a SIRT3 activator [22], on the human SIRT3 enzyme under equilibrium, steady state and non-steady state kinetic conditions. We elucidate the mechanism whereby honokiol activates SIRT3 under physiologically relevant conditions and establish the principles whereby hit compounds like honokiol can be evolved into more potent SIRT3 activator lead compounds.

## MATERIALS AND METHODS

### Chemicals and Reagents

MnSOD (KGELLEAI-(KAc)-RDFGSFDKF) was synthesized by GenScript (Piscataway, NJ). FdL2 (QPKK^AC^-AMC) peptide, also called p53-AMC peptide, was purchased from Enzo Life Sciences (Farmingdale, NY). carba-NAD (Fig. S1) was synthesized by Dalton Pharma (Toronto, ON). Honokiol (HKL) was purchased from Sigma (St. Louis, MO). The solubility profiles of these compounds were determined using HPLC. All other chemicals, solvents used were of the highest purity commercially available and were purchased from either Sigma (St. Louis, MO) or Fisher Scientific (Pittsburgh, PA) or from VWR International (Bridgeport, NJ). The plasmid containing human Sirt3^102-399^ (pEX-His-hSIRT3^102-399^) was purchased from OriGene (Rockville, MD).

### Expression and purification of hSirt3^102-399^

The hSirt3^102-399^ was expressed in *E.coli* Arctic Express (DE3) cells (Agilent Technologies, Wilmington, DE) as per manufacturer’s recommendations. A single bacterial colony was inoculated in LB media containing ampicillin and gentamycin at 37°C. For protein purification, we inoculated overnight grown culture into 200 ml LB medium and grown at 30°C, 250 rpm for 4 hours. We induced Sirt3 expression by adding 1 mM IPTG at 15°C. After 24 hours, cells were harvested, and the pellet was re-suspended in A1 buffer (50 mM NaH_2_PO_4_, 300 mM NaCl, 10 mM imidazole, pH 8.0) and sonicated. A clear supernatant was obtained by centrifugation at 14000 x g for 25 min at 4°C then loaded onto a 5 ml HisTrap HP column, pre-equilibrated with A1 buffer, attached to an AKTA pure FPLC system (GE Healthcare, Pittsburgh, PA). The column was then washed with 50 ml each buffer A1, A2 (50 mM NaH_2_PO_4_, 300 mM NaCl, 75 mM imidazole, pH 8.0), A3 (20 mM Tris-HCl, 2M urea, pH 6.8), and again with buffer A2. Protein was eluted with buffer B1 (50 mM NaH_2_PO_4_, 300 mM NaCl, 300 mM imidazole, pH 8.0). The protein fractions were pooled, dialyzed against dialysis buffer (25 mM Tris, 100 mM NaCl, 5 mM DTT, 10% glycerol, pH 7.5) at 4°C overnight. The purity of final protein was >85% as assessed by SDS-PAGE.

### hSirt3^102-399^-ligand binding assay by Microscale Thermophoresis

DY-647P1-NHS-Ester conjugated hSirt3^102-399^ was used for the MST studies. Briefly, the protein conjugation was done in 1x PBS pH 7.4, 0.05 % Pluronic F-127, and the labeled protein was buffer exchanged into 47 mM Tris-HCl pH 8.0, 129 mM NaCl, 2.5 mM KCl, 0.94 mM MgCl_2_, 5% DMSO, 0.05% Tween-20. We used 2 nM labeled hSirt3 and titrated with varying concentrations of the modulators in the absence and presence of various concentrations of substrates, products and intermediates. The thermophoresis was measured (excitation/emission 653/672 nm) using a Monolith NT. 115 Pico (NanoTemper Technologies) at 25°C. Dissociation constants (*K*_*d*_) were determined using GraFit7 (Erithacus Software) by nonlinear fitting.

### Effect of Honokiol on hSirt3^102-399^ deacetylation activity

Enzymatic reactions included either 2.5 mM NAD^+^ and 6.25 µM MnSOD peptide or 50 µM NAD^+^ and 600 µM peptide substrate in presence of HKL (range 0-200 µM), in a buffer (50mM TRIS-HCl, 137mM NaCl, 2.7mM KCl, and 1mM MgCl_2_, pH 8.0 and 5% DMSO). We started the reactions by adding 5U (0.194 μM) hSirt3^102-399^ and incubated at 37°C for 30 minutes. We terminated the reactions by TFA. The peptide product and substrate were resolved using HPLC; see Supporting Information (SI) for the detailed method.

### hSirt3^102-399^ kinetic analysis using Fluorolabeled Peptide

We determined the steady state kinetic parameters of hSirt3^102-399^ deacetylation using FdL2 peptide. The enzymatic reactions were carried out similar to as described above. For initial velocity determination, we performed the reaction at different [NAD^+^] and in presence of 100 µM FdL2 peptide. We terminated the reactions at specified time by adding the 1X developer and measured the fluorescence on TECAN microplate reader. The raw data were fitted to the defined model equations using GraphPad Prism (GraphPad Software, Inc, CA).

### Preparation of SIRT3.Ac-Pr.NAD^+^ substrate complex for protein-ligand docking

#### Modeling the SIRT3 complex structure

The SIRT3 structure is prepared by aligning the two crystallographic models available in RCSB-PDB. The structure of protein was chosen from 4BVG and the NAD and peptide are chosen from 4FVT. The ligand exchanges Carba-NAD into NAD^+^. The Complex structure was treated with Protein Preparation Wizard module of Schrdinger suite. The module assigns the bond orders and adds hydrogens based on Epik calculations for proper pKa values. Het molecules except for native ligands were eliminated and optimized of their complex structure by Prime module of Schrodinger suite. The side chain structure of HIS248 has been modified. The histidine was defined HIE, because N of the histidine is closed oxygen of ribose of NAD.

The receptor was further equilibrated by MD simulations using the Desmond package prior to QM/MM geometry optimization as described in Supporting Information.

#### QM-MM optimization of the SIRT3 complex

The minimized SIRT3-ALY-NAD^+^ complex structure optimized using the OPLS2005 force field was further optimized at the Q-site (DFT-B3LYP/6-31G*: OPLS2005) level. The quantum mechanics (QM) region of reactant includes the NAD^+^, acetyl lysine entire residue of peptide, GLN 228 entire residue of receptor and side chain (CH2C3N2H3) of HIS248, for a total of 132 atoms. The QM region was calculated by using the density functional theory with the B3LYP exchange-correlation functional and 6-31G* basis set. The remainder of the system (MM region) was treated by using the OPLS2005 force field. A total of 4023 atoms in the system were included for the QM/MM simulations by using the Qsite module as implemented in SCHRODINGER. To avoid over polarization of QM atoms by MM atoms to the boundary, Gaussian charge distributions were used to represent the potential of the atoms within two covalent bonds of the QM/MM cut-site using the Gaussian grid method for hydrogen cap electrostatics in QSite. MM point charges were employed for the rest of the MM region. A total of 4023 atoms were included for the QM/MM calculations by using the Qsite module in SCHRODINGER.

### Protein-ligand docking

Molecular dockings of compounds to the QM/MM optimized structure of SIRT3:Ar-Pr-NAD^+^ substrate complex have been carried out by AutoDockVina. AutoDockVina repeatedly docks each ligand to the target applying several search algorithms like global search implementing simulated annealing genetic algorithm, local search algorithm hybrid globallocal search algorithm and also Lamarckian genetic algorithm and iterated local search, i.e. genetic algorithm with local gradient. AutoDockTools have been used to prepare the receptor PBQT format adding polar hydrogen and assigning Gasteiger charges to all its atoms as well as identifying the coordinates of the target box to enclose QM/MM optimized SIRT3.Ac-Pr.NAD^+^ substrate for global docking with a cubic grid box of 126×126×126 point dimensions of 0.5 resolution. Prior to docking, Honokiol structure was geometry optimized in gas phase by Hartree-Fock method with 6-311G+ basis.

### Loop modeling

The following protocol was used to prepare starting structures for molecular dynamics simulation of the SIRT3 ternary (SIRT3+AcPr+NAD^+^), intermediate (SIRT3+ADPR-Pr-Im) and product (SIRT3+ Pr +AADPR) complexes with native and nonnative loop. For the ternary complex (prepared based on the 4FVT crystal structure for all protein residues except the cofactor binding loop), the intermediate (closed) conformation of the cofactor binding loop (residues 155-178) was substituted, whereas for the intermediate complex (prepared based on the 4BVG crystal structure), the ternary (open) conformation of the cofactor binding loop (residues 155-178) was substituted. For the coproduct complex (prepared based on 4BVH crystal structure), the apo (open) conformation of the cofactor binding loop (residues 155-178 from the 3GLS crystal structure) was substituted. The following steps were applied:

- Sequence-structure based superimposition
- Graft the coordinates for the co-factor loop region.
- Run protein preparation wizard on the modelled product complex (open/closed loop conformation).
- Predict/repack the side chains for all residues within 7.5 Å of the grafted loop region using Monte Carlo approach together with backbone sampling.
- Carry out prime energy refinement only on those residues which were repacked keeping the others fixed.

### Molecular dynamics simulation

#### Force field

The Amber99SB force field [28, 29] was used for all the molecular mechanics calculations. Extra parameters were adapted from the following sources: parameters for Zn developed by Luo’s group [30]; parameters for NAD^+^ developed by Walker et al[31] and Pavelites et al [32] ; parameters for acetylated lysine developed by Papamokos et al [33]. Amber14 tools (Antechamber program) are employed for parametrizing the non-standard residues and tleap program for creating topology and force field parameters for the standard residues.

#### Fast long-range electrostatics

The Particle Mesh Ewald (PME) method [34] was used to calculate the electrostatic energy.

#### Explicit vs implicit solvation

Explicit solvent for sampling but implicit was used for more energy calculations. Water molecules were stripped from the molecular dynamics trajectories to prepare for MM/PBSA and MM/GBSA re-runs.

#### MD simulation

All molecular dynamics simulations were performed with periodic boundary conditions to produce isothermal-isobaric ensembles (NPT) at 300 K using the NAMD program [35]. The integration of the equations of motion was conducted at a time step of 2 femtoseconds. The covalent bonds involving hydrogen atoms were frozen with the SHAKE algorithm [36]. Temperature was regulated using the Langevin dynamics with the collision frequency of 1 ps^-1^. Pressure regulation was achieved with isotropic position scaling and the pressure relaxation time was set to 1.0 picosecond.

Conformational energies were calculated by the MD/MM-GB(PB)SA method, as described in Supporting Information.

Additional methods employed are described in the Supporting Information (SI).

## RESULTS

The sirtuin catalytic cycle (Fig. 1) is believed to proceed in two consecutive stages [2]. The initial stage (ADP-ribosylation) involves the cleavage of the NAM moiety of NAD^+^ and the nucleophilic attack of the acyl-Lys side chain of the protein substrate to form a positively charged O-alkylimidate intermediate [2, 7]. NAM-induced reversal of the intermediate (the so-called base exchange reaction) causes reformation of NAD^+^ and acyl-Lys protein. The energetics of this reversible reaction affects both the potency of NAM inhibition of sirtuins and the Michaelis constant for NAD^+^ (*K*_*m,NAD+*_). The second stage of sirtuin catalysis, which includes the rate-determining step, involves four successive steps that culminate in deacylation of the Lys side chain of the protein substrate and the formation of O-acetyl ADP ribose coproduct [2, 4, 37].

As described in [27] the initial rate of deacylation can be expressed

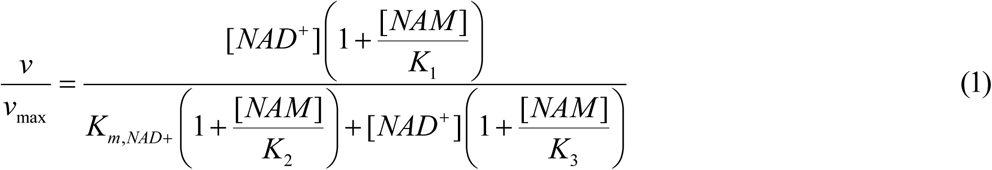

where the definitions of the steady state constants in terms of fundamental rate constants in the enzymatic reaction mechanism are presented in Fig. S2. Expression (1) can be used to calculate the initial rate of sirtuin-catalyzed deacylation for specified intracellular concentrations of NAD^+^ and NAM, assuming the rate constants are known. At constant [NAM], Eq. (1) is typically represented graphically as double reciprocal plots, where the slope of the plot (1/v *vs* 1/ [NAD^+^]) at [NAM] = 0 is *K*_*m,NAD*+_ / *v*_max_, for which the expression is:

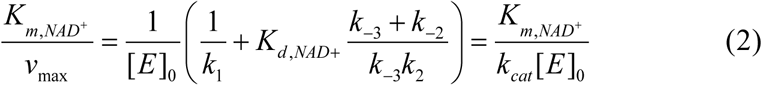

The steady state parameter *α*, which is a measure of the extent of competitive inhibition by the endogenous inhibitor NAM against the cofactor NAD^+^, can be expressed in terms of the ratio of *K*_*d,NAD+*_ and *K*_*m,NAD+*_ [38]:

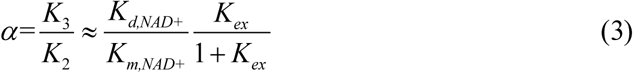

where *K*_*ex*_ = k _-2_/k_2_ and the approximations are defined in Fig. S2.

For enzyme activation to be possible, the modulator must co-bind with substrates – namely, NAD^+^ and acylated peptide in the case of sirtuins. Within the context of traditional enzyme inhibition, two modes of action satisfy this condition: noncompetitive and uncompetitive inhibition. Examples of noncompetitive inhibitors of sirtuins include SirReal2 [39], whereas examples of uncompetitive inhibitors include Ex-527 [40]. Although some such sirtuin inhibitors may satisfy the requirement of cobinding with substrates, they do not possess other critical attributes needed for mechanism-based enzyme activation. While such compounds may have promising properties as potential hits for the development of mechanism-based activators, most previous work has only characterized their kinetic effects using traditional rapid equilibrium enzyme inhibition formulations, rather than a steady-state formulation for mechanism-based enzyme modulation. In our previous study [27], we presented a theory of mechanism-based enzyme activation, aimed at establishing a foundation for the characterization and design of nonallosteric enzyme activating compounds. Fig. S3 illustrates the theoretical model for mechanism-based sirtuin enzyme activation schematically. This theory provides a unifying framework under which such compounds can be characterized and hits can be evolved into leads. Here, we apply this theory to experimental data we collected for reported nonallosteric activators of sirtuins.

Appendix A1 provides the expressions for each of the modulated steady state constants in Fig. S2 in the presence of a specified concentration [A] of the hit compound, according to the mechanism-based enzyme activation theory presented in [27]. The *K*_*di,A*_ in Appendix A1 denote the dissociation constants of the hit compound with respect to the respective reaction intermediates indexed by *i* in the model schematic Fig. S3.

Fig. 2 depicts the model-predicted changes to the various steady state, Michaelis and dissociation constants in the sirtuin reaction mechanism in the presence of such a modulator.

**Fig. 2.**
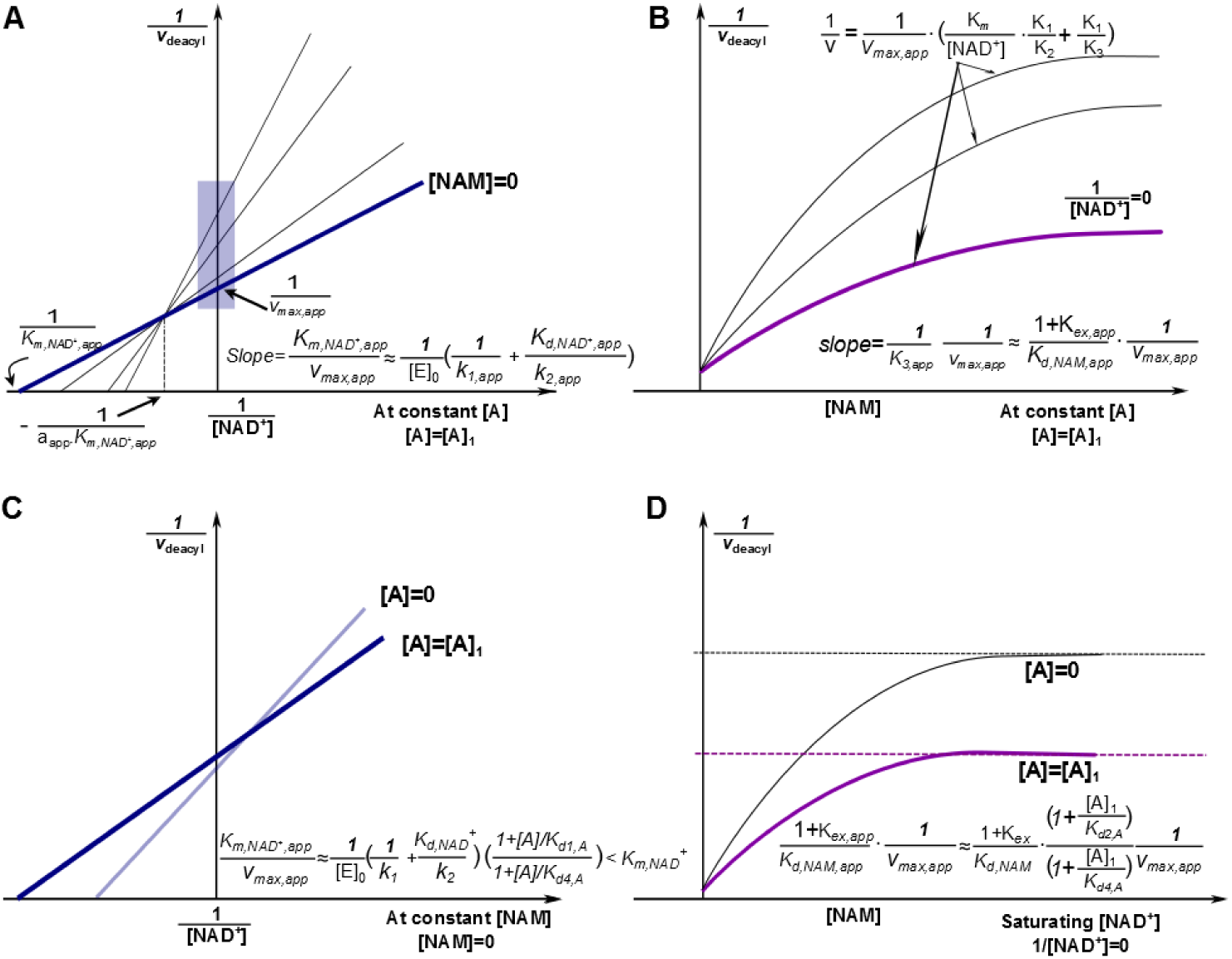
Mechanism-based activation of sirtuin enzymes: predicted steady-state properties and dose-response behavior. **(A)** Double reciprocal plots for deacylation initial rate measurements in the presence of activator. The blue box on the y-axis highlights the data that is used to construct the Dixon plot at saturating [NAD^+^] depicted in (B). **(B)** Dixon plots for deacylation initial rate measurements in the presence of activator. The arrows point to predicted plateaus in these curves. **(C)** Comparison of double reciprocal plots at [NAM] = 0 µM in the presence and absence of activator. **(D)** Comparison of Dixon plots at 1/[NAD^+^] = 0 in the presence and absence of activator. “A” denotes a mechanism-based sirtuin activating compound.

Note that the model depicted omits the term quadratic in [NAM] in Eq. (S1) in SI and the plateaus/dotted lines shown in the Dixon plots are the asymptotic values to which the model-predicted rates converge in the absence of this term. Within this model, the conditions for activation were described in [27]. These conditions establish the biophysical properties of the compound that are required to elicit changes in the steady state constants that are conducive to activation. A hit compound may be defined as one that satisfies a subset of the conditions enumerated, and may also display comparatively little inhibition in the pre-steady state burst phase. For instance, it is possible that a hit compound increases catalytic efficiency in the presence of NAM (for example, due to increase in the *K*_*3*_ parameter above), but not in its absence. It may be possible to further improve these compounds to induce substrate-specific activation of sirtuins like SIRT3, under physiologically relevant NAD^+^ depletion conditions. Note that because there is a nonzero physiological concentration of NAM in the cell, reducing NAM inhibition can also contribute to activation under conditions that are relevant physiologically. Alternatively, the relative rates of deacylation in the presence and absence of modulator could converge under certain combinations of [NAD^+^] and [NAM].

### Computational characterization of reported hit compounds for mechanism-based sirtuin activation

Typically, mechanism-based activators [27] may alter the distributions of local degrees of freedom whose conformations change during the course of the reaction, which leads to tradeoffs in the ΔΔG’s for various reactions steps upon stabilization of one such conformation (see below). This is a critical distinction from allosteric activation, where there is no conformational change specific to certain steps in the reaction.

Activators of sirtuin enzymes that do not possess an allosteric site [41, 42] have been shown crystallographically to induce changes in the conformation of the so-called flexible cofactor binding loop in sirtuin enzymes. This loop changes conformation after the first chemical step of the reaction, namely the cleavage of nicotinamide from the NAD^+^ cofactor. In these cases, the local degrees of freedom above are the backbone degrees of freedom in the flexible cofactor binding loop. Fig. 3 displays a structure alignment illustrating this loop conformational change that occurs during the catalytic cycle of sirtuin enzymes.

**Fig. 3.**
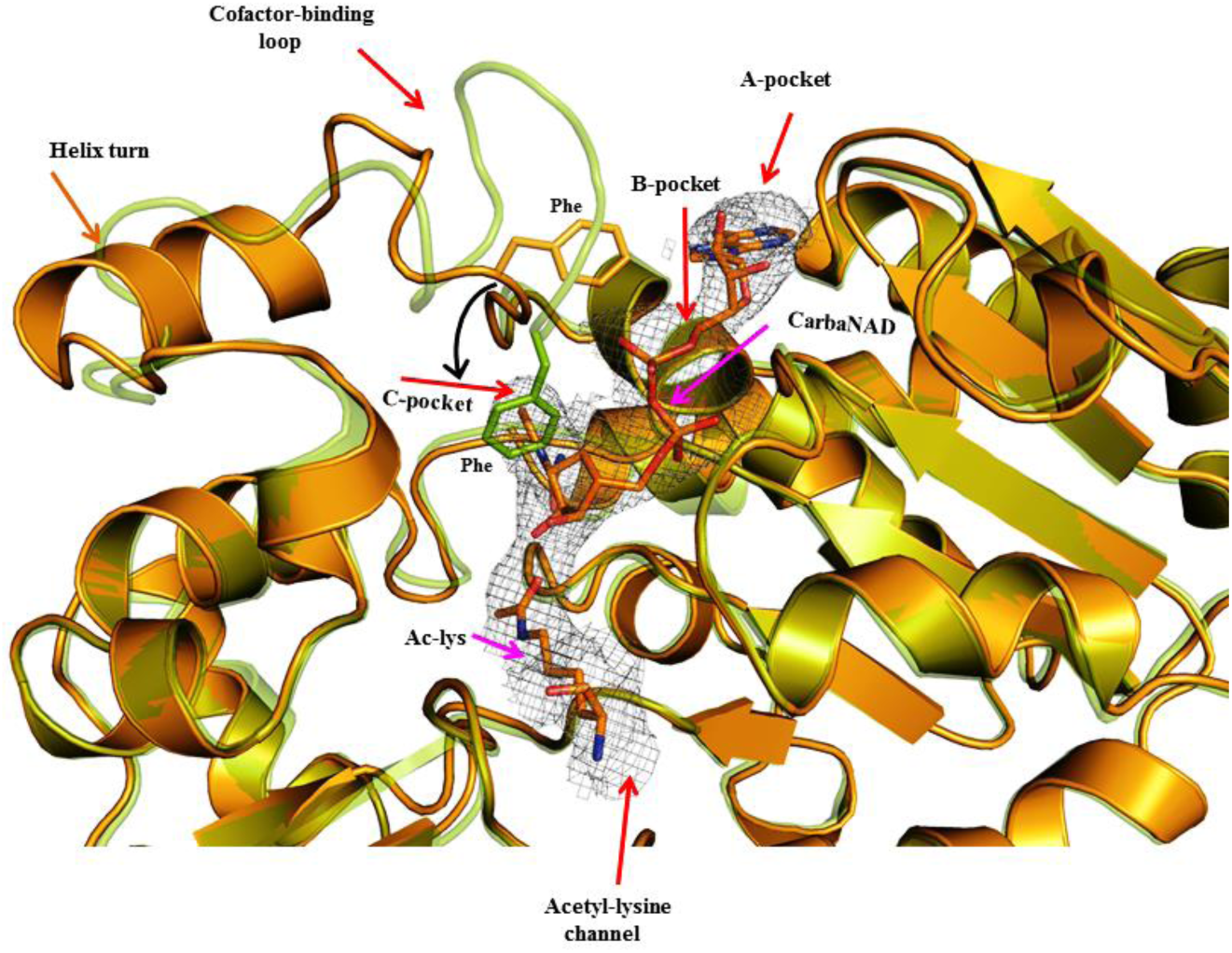
Superposition of Sirt3 native intermediate (4BVG - Green) and Sirt3 ternary complex (4FVT - Orange) showing differences in the conformation of the cofactor binding loop and the position of the Phe residue. Individual subsites are highlighted and the movement of Phe residue is indicated by black arrows. The substrates Carba-NAD and Ac-Lysine are rendered in stick representation.

Along with SIRT1, SIRT3 is one of the most important sirtuin enzymes involved in the regulation of healthspan [43]; hence we chose it as a subject of study. Molecular dynamics simulations were employed to study the flexibility of the SIRT3 cofactor binding loop in different conformations associated with the stages of the sirtuin catalytic cycle (Fig. 4). The high B factors for the ternary complex cofactor binding loop are consistent with the ability of modulator binding near the loop to alter its conformation.

**Fig. 4.**
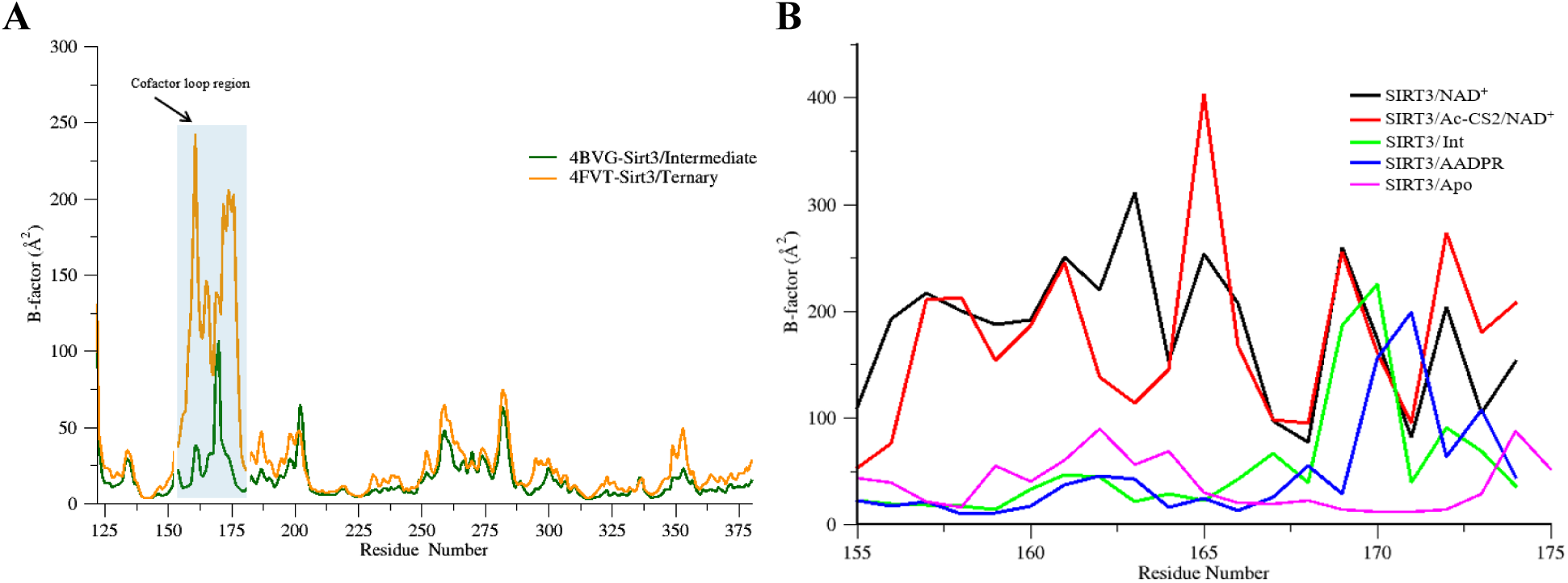
Plot showing simulated B-factor values for C_α_ atoms belonging to (A) residues 125-375; (B) the co-factor binding loop region of various SIRT3 complexes. Residues (162-170) are known to adopt a helix conformation when bound to substrate. Ternary = Ac-CS2/NAD^+^ complex, Int=Intermediate

Honokiol (HKL) (Fig. 8C) has been reported as a SIRT3 activator for the MnSOD protein substrate [22]. Because SIRT3 does not share the allosteric activation site of SIRT1, we studied HKL as a potential hit compound for mechanism-based activation of SIRT3. The binding mode of HKL to human SIRT3 was investigated computationally through protein-ligand docking to the SIRT3:NAD^+^:AcPr ternary reactants complex (Fig. 5), which demonstrated high affinity (−5.8 kcal/mol) co-binding of HKL to a pocket above the active site. Honokiol is predicted to engage in H-bonds with Glu177 of the flexible cofactor binding loop which will induce structural changes due to loop flexibility. As such, we carried out molecular dynamics simulations for two different conformations of the cofactor binding loop in the ternary reactants complex – and open and closed loop conformation, and compared the interactions between the loop and NAD^+^ in these two conformations (Fig. 6). Whereas Arg 38 engages in favorable electrostatic interactions with NAD^+^ upon loop closure, Phe 158 (not depicted) induces strain in the nicotinamide moiety of NAD^+^, which typically prevents loop closure in the absence of a modulator.

**Fig. 5.**
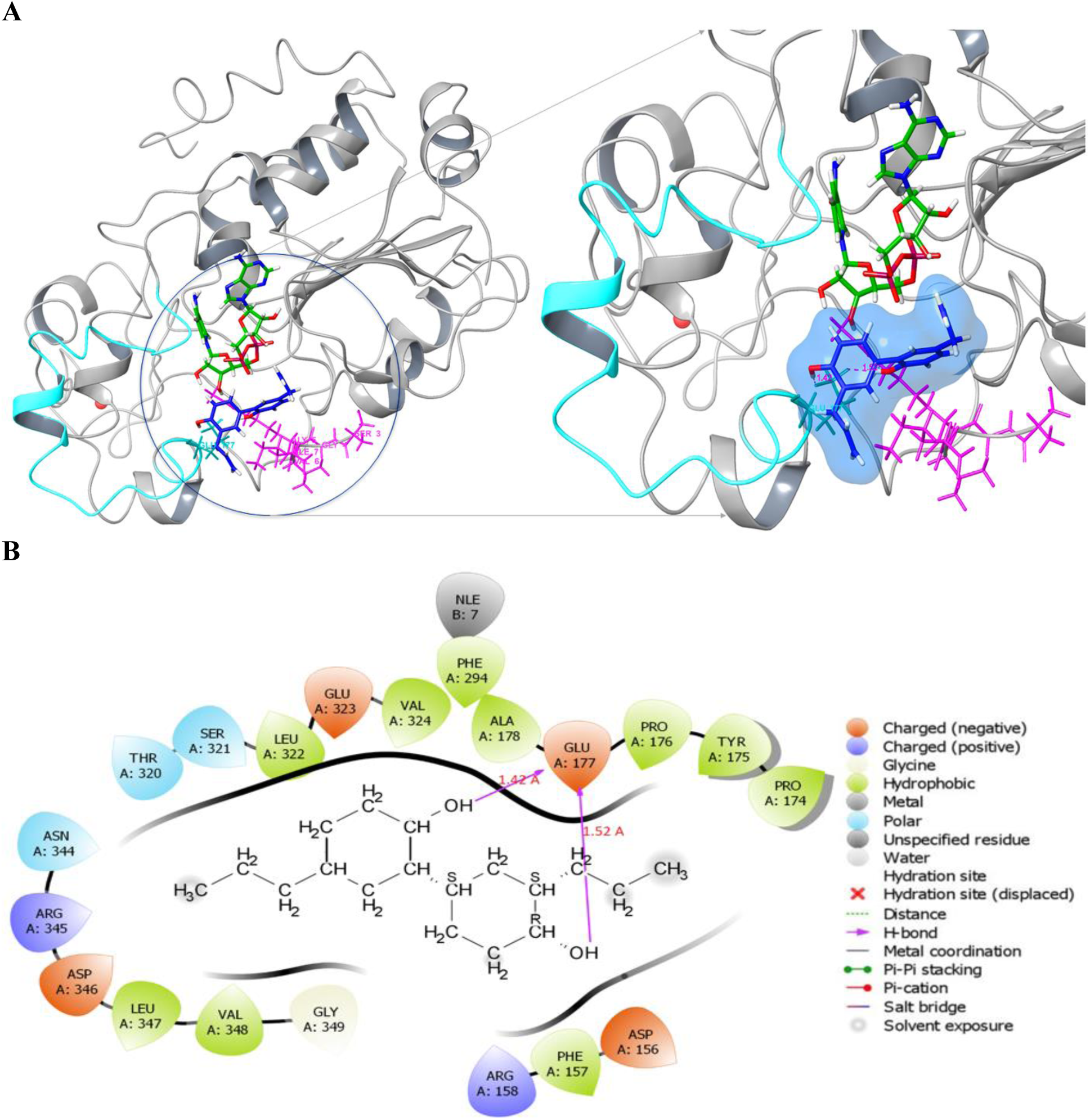
(A) Binding mode of Honokiol to the SIRT3 ternary complex (SIRT3: Ac-ACS2: NAD^+^) by docking. The SIRT3 protein is represented in its secondary structure colored in white. NAD^+^ in green, Ac-ACS2 in pink, Honokiol in blue. The Honokiol molecule forms H-bonds with Glu 177 which is present in the co-factor binding loop (in cyan) of the SIRT3 protein. (**B) The ligand interaction diagram of Honokiol in the SIRT3 ternary complex (SIRT3: Ac-ACS2:NAD**^**+**^). The Honokiol ligand binds with the SIRT3 protein at Glu 177 position and forms two hydrogen bonds with two of its hydroxyl groups present in it. The Glu 177 is present in the cofactor binding loop of SIRT3 protein.

**Fig. 6.**
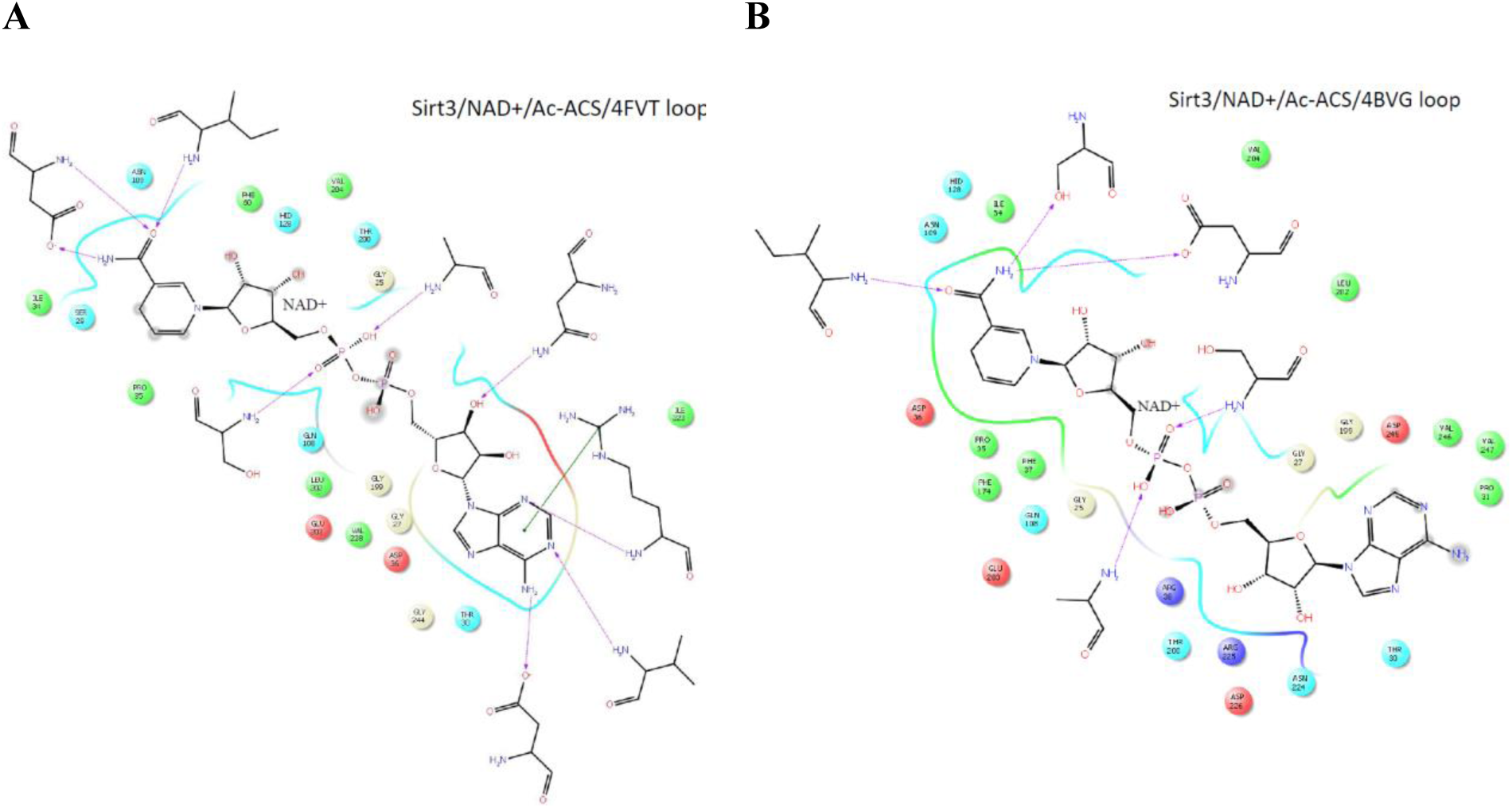
NAD^+^ ligand interaction diagrams for the SIRT ternary reactants complex with (A) open and (B) closed loop conformations depicting the significant changes in the residue interactions with NAD^+^ upon closing of the loop induced by modulators. Geometries depicted are MD averages.

**Fig. 7.**
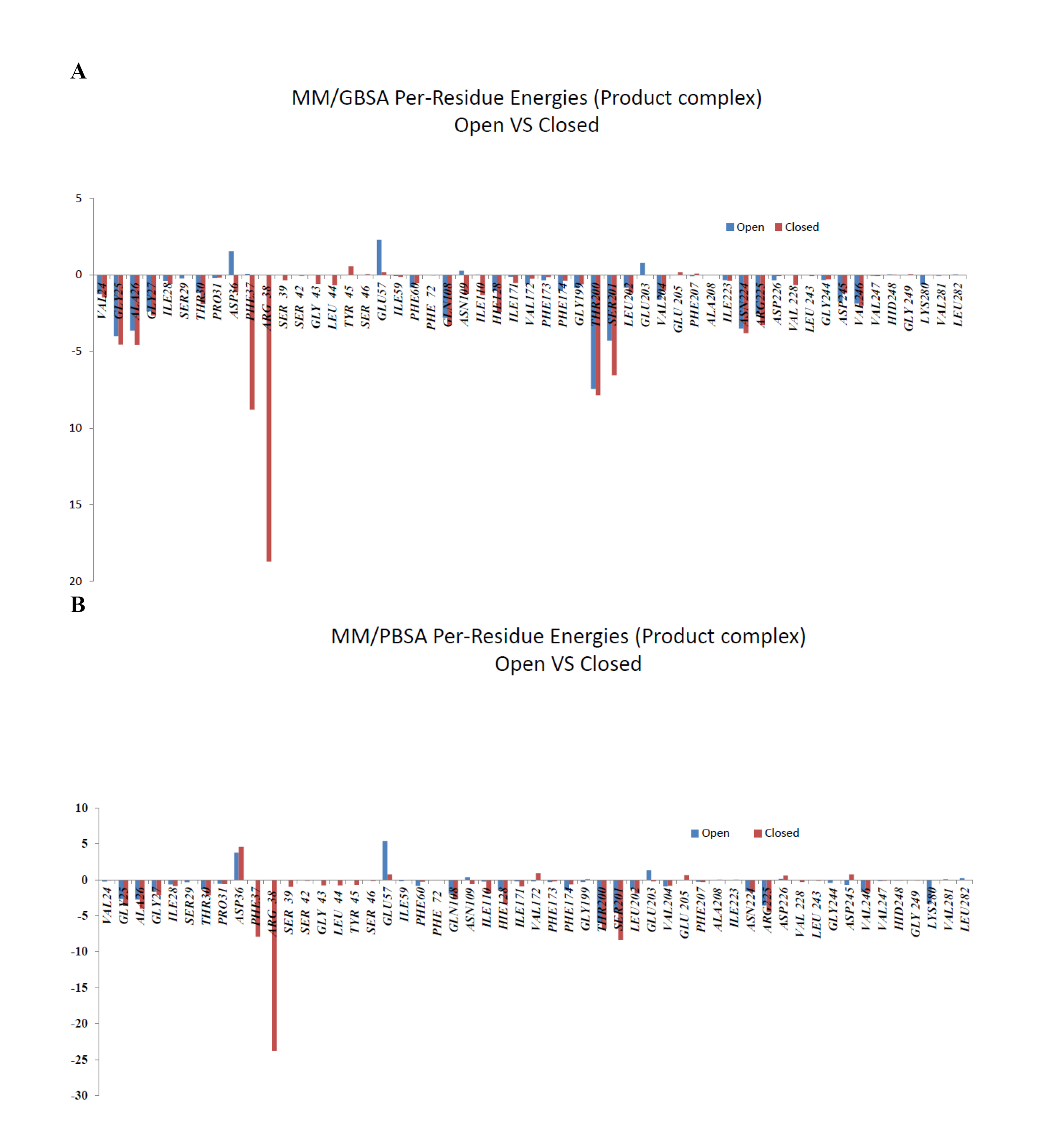
By-residue (A) MM-GBSA binding energies and B) MM-PBSA MD average binding affinities of SIRT3 to the AADPR coproduct for open vs closed loop conformations.

**Fig. 8.**
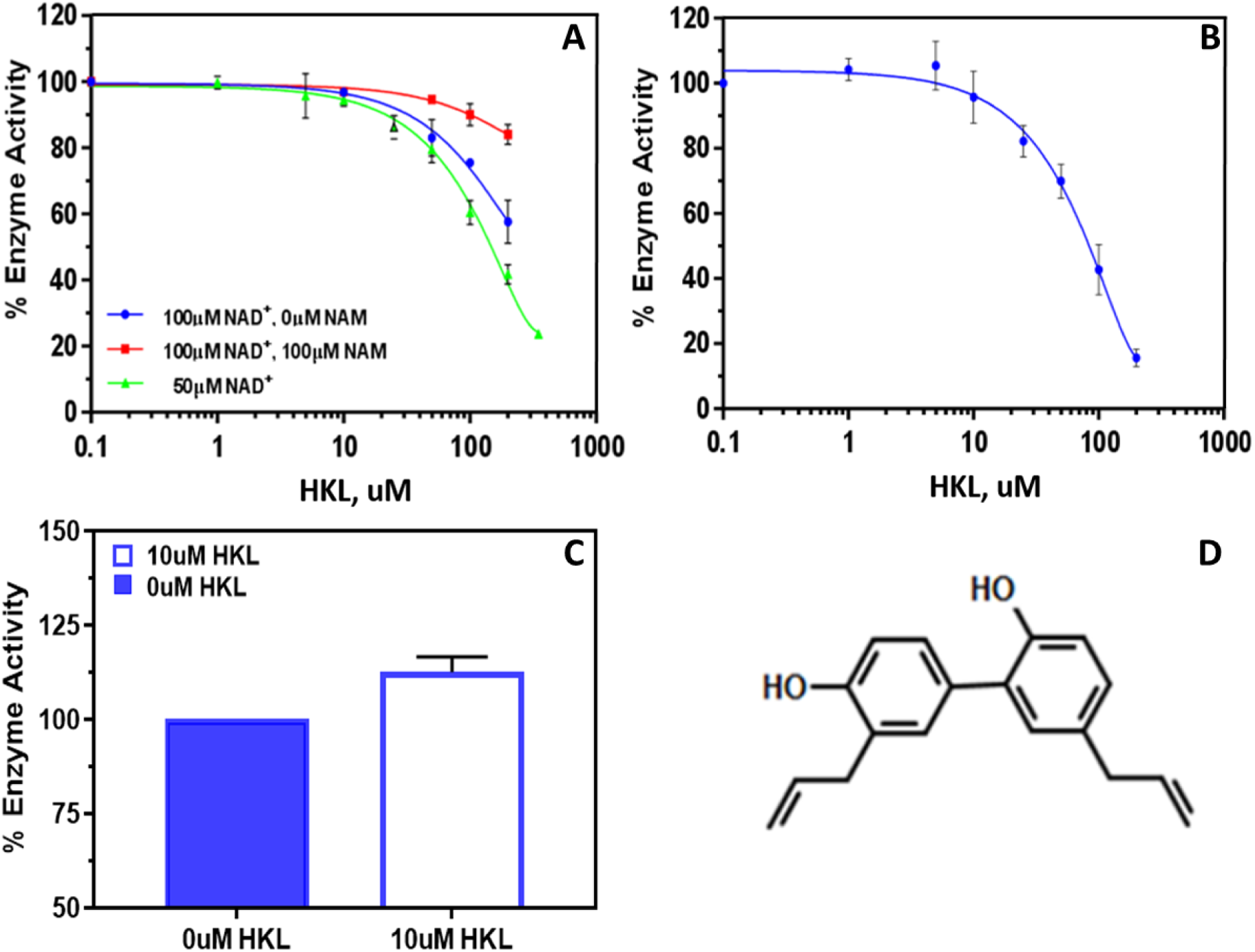
Effect of HKL on Sirt3 deacetylation activity using a label-free assay: dose-response. Dose-response curves were measured under conditions where [E]_0_/[S]_0_ << 1, where [S]_0_ denotes the initial concentration of the limiting substrate, in order to maximize the contribution the steady state phase of the reaction to the curve. **(A)** The superposition of the un-saturating NAD^+^ and saturating MnSOD K122 peptide in the three formats of (blue) 10 mins, 100 µM NAD^+^ (N=2); (red) 10 mins, 100 µM NAD^+^/100 µM NAM (N=2), and (green) 30 min, 50 µM NAD^+^ (N=3). **(B)** 2.5 mM NAD^+^ and 6.25 µM un-saturating MnSOD K122 peptide (N=5). **(C)** Activation of Sirt3 in the presence of 10 uM HKL at [E]_0_/[NAD^+^]=0.0176 ([E]_0_=0.8807uM, [NAD^+^]=50 uM, [MnSOD]=600uM, 1.5 min, N=6). **(D)** Structure of HKL, 5, 3’-Diallyl-2, 4’-dihydroxybiphenyl. The lines are the results of global fitting; the lines were not fit independently to the data points in the individual panels. The error bars that are not visible are too small to view at this scale.

In addition, in order to investigate the effect of HKL-induced changes of the cofactor binding loop conformation on coproduct binding affinity, we calculated O-AADPR binding affinities from molecular dynamics simulations for open and closed conformations of the loop (Fig. 7). The significantly higher binding energies for the closed loop conformation demonstrate that stabilization of the closed loop conformation will increase coproduct binding affinity and reduce the rate of coproduct dissociation.

### Experimental characterization of proposed hit compounds for mechanism-based sirtuin activation: kinetic studies

In this section we experimentally characterize proposed sirtuin-activating compounds within the context of the mechanistic model above, evaluating their characteristics within the context of the ideal features of mechanism-based activators. Both thermodynamic and kinetic measurements are made in order to characterize the equilibrium and steady state constants entering the mechanism-based enzyme activation model. We begin with the kinetic measurements.

Because such mechanism-based modulators operate according to new modes of action, as depicted in the model schematic Fig. S3, the traditional approach of characterizing a compound as uncompetitive, noncompetitive or competitive with respect to substrates is not sufficient, even if the modulator has a net inhibitory effect. The model introduced in previous study [27] and summarized here provides general principles for kinetic characterization of mechanism-based enzyme activating compounds, which we apply below.

The aim of steady state characterization of hit compounds for mechanism-based activation is to estimate the parameters in Eq. (1) in the absence of modulator and at a saturating concentration of modulator, varying both [NAD^+^] and [NAM] in each case. This provides estimates of both the front and back face steady state parameters in the model schematic Fig. S3A. By contrast, a mixed inhibition model with respect to the modulator concentration (as, e.g., applied to certain mechanism-based inhibitors [39]) would not have an interpretation for the steady state constants in terms of fundamental rate constants in the sirtuin mechanism, whereas characterizing unsaturating modulator concentrations at each of multiple product (NAM) concentrations, while providing more information, would similarly estimate only apparent steady state parameters depicted in Fig. S3B.

Experimental characterization of HKL on SIRT3 activity was carried out according to the principles of mechanism-based enzyme activation. Dose-response curves are measured at [E]_0_/[S]_0_ <<1, where [S]_0_ is the initial concentration of the limiting substrate (NAD^+^ or peptide) (Fig. 8A, B). Fig. 8A and C illustrate how for [E]_0_/[S]0 << 1, which is required for steady state characterization, where S is the limiting substrate, the measurement time has a significant impact on the effect of HKL on enzyme activity, with activation occurring at short times (pre-steady state). Both the green curve in Fig. 8A and the bar plot in Fig. 8C were assayed at 50uM NAD^+^, but at significantly different times (30 min and 1.5 min, respectively). The red curve in Fig. 8A and the bar plot in Fig. 8C were assayed at progressively shorter times (10 min and 1.5 min respectively), but at different concentrations of NAD^+^ and NAM, which affect the duration of the pre-steady state phase of the reaction. The observed activation of SIRT3 by HKL in Fig. 8C highly statistically significant, with p < 0.001.

Fig. S10 depicts how HKL also activates SIRT3 under non-steady state conditions for the p53-AMC peptide substrate, analogously to the results in Fig. 8C for the MnSOD substrate. This fluorolabeled substrate can be employed using the reaction conditions depicted in this Figure to rapidly screen for hit compounds that nonallosterically activate SIRT3 due to the sensitivity fluorescence-based activity assays. Activity assays with the fluorolabeled p53 substrate were repeated with HPLC (Fig. S10) to eliminate the possibility of false positives.

The steady state rate measurements at multiple values of [NAD^+^] and [NAM] were carried out in order to estimate the parameters in model eq. (2). The results for the MnSOD substrate are presented in Figs 9, 10, and Table 1. Note that at very high values of [NAM], the effect of differences in *K*_*d,NAM*_ induced by the modulator becomes negligible, allowing isolation of differences in the NAD^+^ binding and ADP ribosylation / base exchange kinetics. Below, we compare the observed changes in the steady state parameters *v*_max_, *K*_*m, NAD*+_, *K*_1_, *K*_2_, *K*_3_ at 200 µM concentration (approximately saturating) to the changes conducive to activation that were delineated above and in previous study [27].

**Fig. 9.**
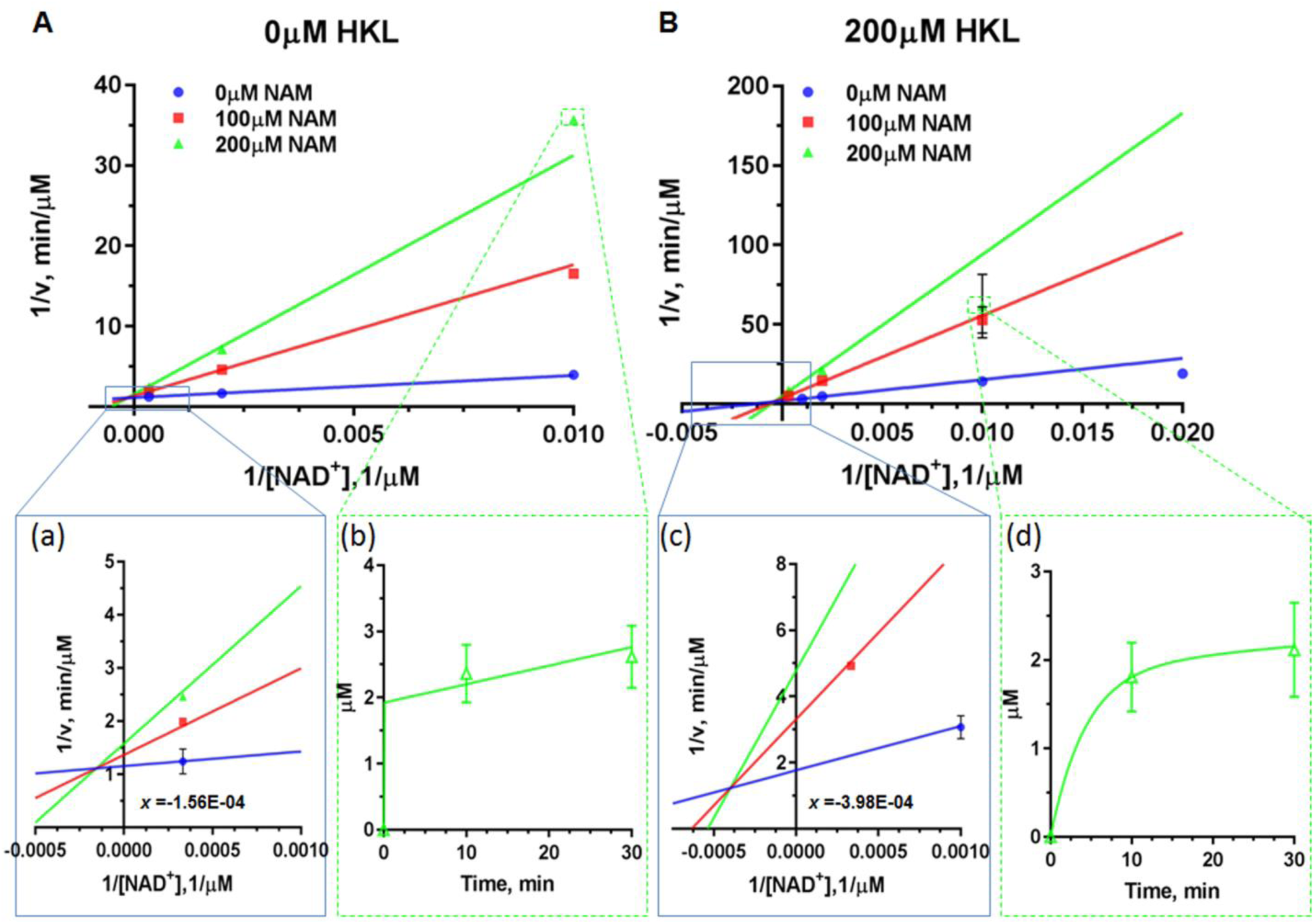
Steady-state kinetic characterization of deacetylation in the presence and absence of the mechanism-based modulator HKL. Double reciprocal plots for deacetylation initial rate measurements of saturating substrate peptide (MnSOD K122) in the presence of different concentrations of NAM with **(A)** 0 μM HKL; **(B)** 200 µM HKL. Enlargements of the intersection points are provided as insets (a) and (c). The time series plots of µM product formed vs. time are provided as insets (b) and (d). The lines are the results of global fitting; the lines were not fit independently to the data points in the individual panels. The error bars that are not visible are too small to view at this scale.

**Fig. 10.**
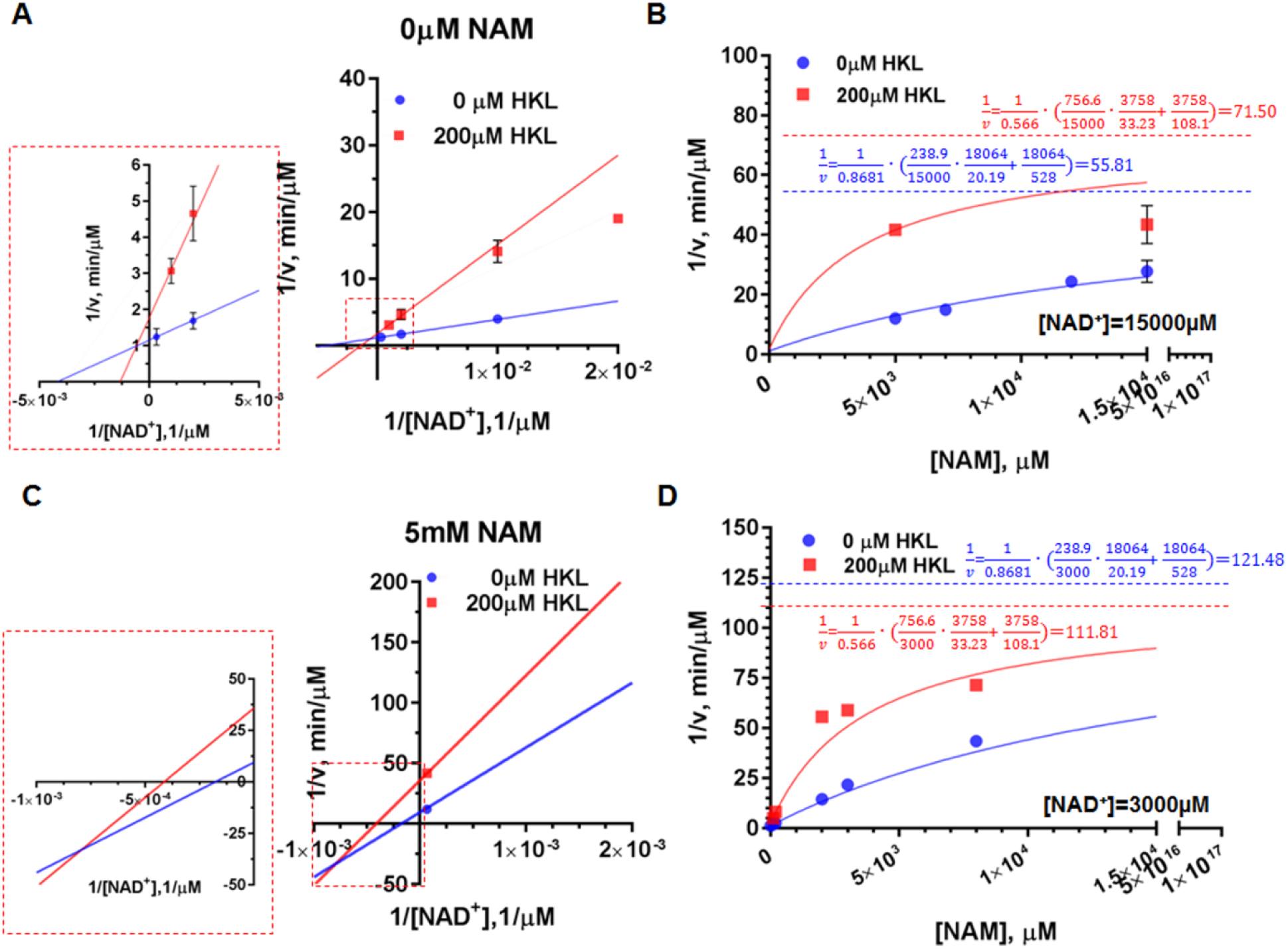
Mechanism-based modulation of hSirt3^102-399^ deacetylation of MnSOD substrate: effect of NAD^+^ and NAM. **(A)** Double reciprocal plots for deacetylation initial rate measurements in the presence and absence of HKL at [NAM] = 0 µM. The enlargement of intersection point is provided as inset. **(B)** Comparison of Dixon plots at [NAD^+^] = 15000 µM in the presence and absence of HKL. **(C)** Double reciprocal plots for deacetylation initial rate measurements in the presence and absence of HKL at [NAM] = 5mM. The enlargement of intersection point is provided as inset. **(D)** Comparison of Dixon plots at [NAD^+^] = 3000 µM in the presence and absence of HKL. Note that the model omits the term quadratic in [NAM] in Eq. (S1) and the dotted lines shown in the Dixon plots are the asymptotic values to which the model-predicted rates converge in the absence of this term.

**Table 1.** Model parameter estimates from global nonlinear fitting of Eq. (1) for SIRT3 in the presence and absence of 200 μM (saturating) HKL. [E_0_] = 1.85 µM.

| | 0$\mu M$ HKL | 200$\mu M$ HKL |
| --- | --- | --- |
| $V_{max}$ | 0.8681 | 0.566 |
| $K_1$ | 18064 | 3758 |
| $K_m$ | 238.9 | 756.6 |
| $K_2$ | 20.19 | 33.23 |
| $K_3$ | 528 | 108.1 |
| **Std. Error** |  |  |
| $V_{max}$ | 0.01216 | 0.05319 |
| $K_1$ | 8279 | 1572 |
| $K_m$ | 12.92 | 138.8 |
| $K_2$ | 1.612 | 6.886 |
| $K_3$ | 105 | 28.98 |
| **95% CI (profile likelihood)** |  |  |
| $V_{max}$ | 0.8417 to 0.8954 | 0.4658 to 0.7421 |
| $K_1$ | 8497 to 288496 | 1826 to 19458 |
| $K_m$ | 212.2 to 268.7 | 501.3 to 1231 |
| $K_2$ | 16.91 to 24.25 | 20.81 to 55.33 |
| $K_3$ | 357.7 to 946.8 | 60.92 to 243.2 |
| **Goodness of Fit** |  |  |
| Degrees of Freedom | 11 | 10 |
| R square | 0.9988 | 0.9947 |
| Absolute Sum of Squares | 0.001052 | 0.0006615 |
| Sy.x | 0.009782 | 0.008133 |

The insets in Fig. 9 show examples of the time series data collected. These insets demonstrate that deacylation of this substrate displays a two phase behavior (pre-steady state/steady state phase) with a relatively slow first phase. Due to this behavior, we used a two-phase rather than one-phase exponential time series fitting. Irrespective of the first phase time constant, the rate in the second phase is almost identical. Hence the uncertainty in steady state rates is very low. The rate is generally almost constant over the measured times in the second phase. Further discussion of time series fitting for the MnSOD substrate are provided in the SI, including Fig. S11.

Because of the apparent two-phase time series dynamics of catalysis for MnSOD, the relative amounts of product formation in the presence vs absence of HKL depends on the choice of measurement time, with a closer correspondence at lower times. In particular, in the presence of 100 µM NAM, which is on the same order of the magnitude as the physiological concentration of NAM, the amounts of product formed at 0 and 200 µM HKL after 10 mins are quite close (Fig. 8A). Compared to the steady state rates, the first phase rates are generally closer in magnitude.

### Measurement of binding affinities

The binding affinities of ligands to complexes in the catalytic mechanism of SIRT3 / MnSOD peptide substrate were measured using microscale thermophoresis (MST; Figs 11, S4). In order to carry out binding affinity measurements on the reactive complex of enzyme, acylated peptide substrate and NAD^+^, we synthesized the catalytically inert molecule carba-NAD^+^ and used this NAD^+^ analog for those MST studies.

**Fig. 11.**
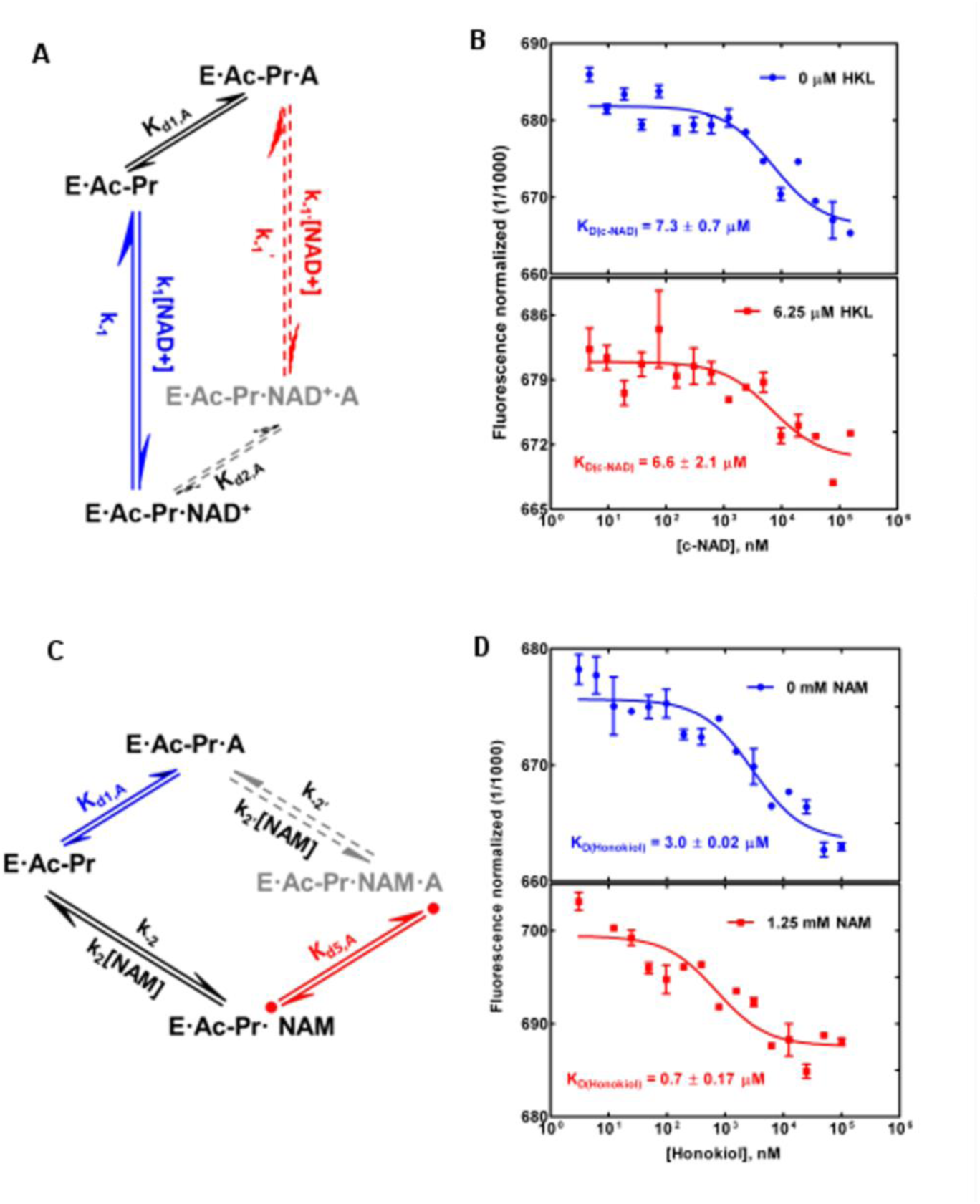
Binding affinity measurements for complexes in the sirtuin reaction mechanism. **(A, C)** Pathways in the sirtuin reaction network from Fig. S3. E, enzyme; Ac-Pr, acetylated peptide substrate; NAD, NAM adenine dinucleotide; A, modulator (HKL); NAM, NAM. **(B)** Carba-NAD binding in the ternary complex: effect of mechanism-based modulator HKL. Binding of carba NAD (c-NAD) to Sirt3.Ac-MnSOD complex, in presence and absence of 6.25 µM HKL measured using MST. **(D)** HKL binding: effect of NAM. Binding of HKL to the Sirt3.Ac-MnSOD complex, in presence and absence of NAM, measured using MST.

In addition to measuring the effect of HKL on NAD^+^ binding affinity, we measured its effect on acylated peptide binding affinity, NAM binding affinity in the presence of acetylated peptide, deacylated peptide binding affinity, O-acetylated ADP ribose binding affinity in the presence of deacylated peptide (i.e., the product complex), and NAD binding affinity in the presence of deacylated peptide. The latter measurements in the presence of deacylated peptide were made in order to determine whether product inhibition plays any role in the observed kinetics. For binding affinity studies on complexes that appear in the model schematic Fig. S3, the relevant faces of the cube are also displayed in the respective figures. Note that measuring the binding affinities for any of the 3 sides of such faces determines the fourth binding affinity. In particular, the binding affinities of HKL or carba-NAD were measured in the ternary complex and product complex. Cooperative binding between HKL and the relevant ligand were studied in each case. It was observed that HKL can have synergistic binding interactions with several ligands.

### Substrate dependence of mechanism-based modulation

The p53-AMC substrate was studied in order to explore the substrate selectivity of modulation by HKL (Fig. 12 and Table S1, Figs. S5 and S6). In enzyme design, the analogous problem of substrate specificity of an engineered protein is often considered. Due to the differing effects of the modulator on these two substrates, which can be identified and characterized within the present framework, substrate selectivity of mechanism-based activation can in principle be engineered.

**Fig. 12.**
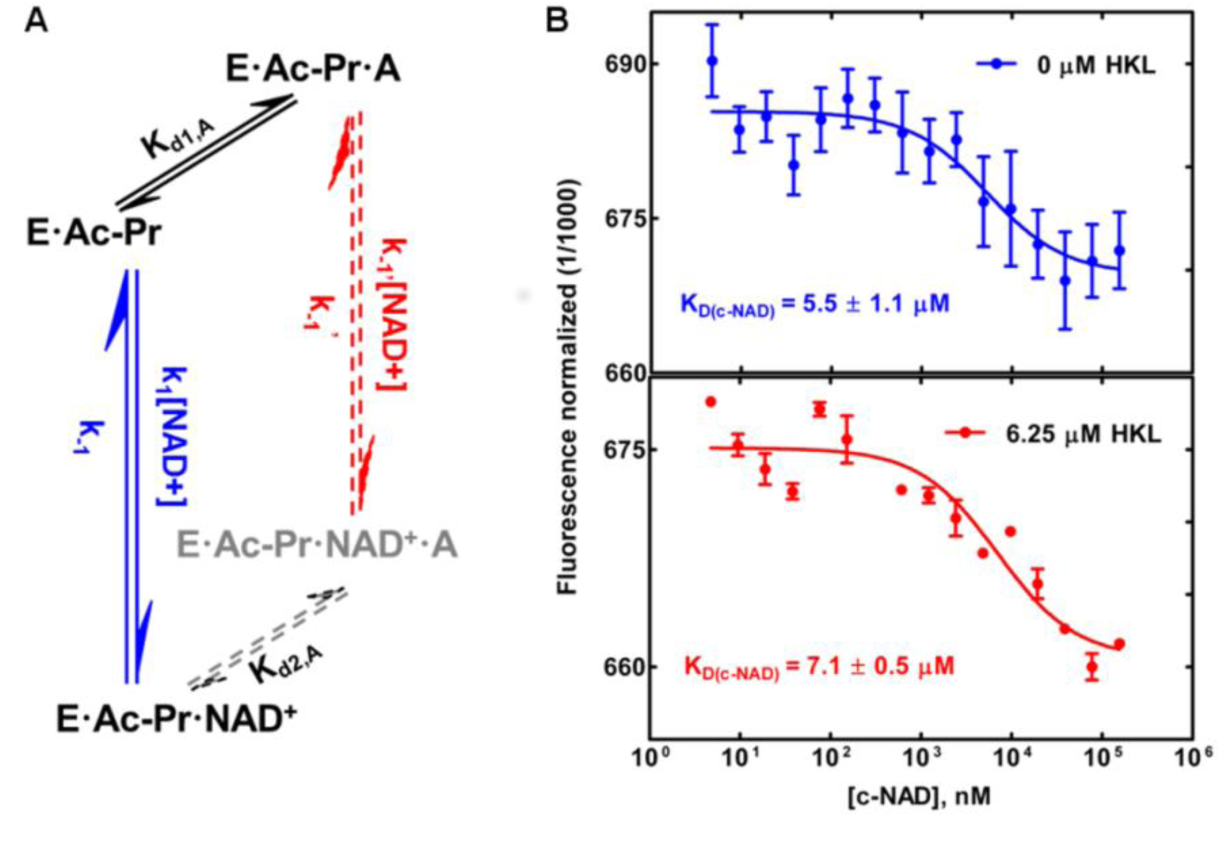
Binding affinity measurements of carba-NAD binding in the ternary complex of an alternate substrate: effect of HKL. **(A)** Pathways in the sirtuin reaction network from Fig. S3. E, enzyme; Ac-Pr, acetylated peptide substrate; NAD, NAM adenine dinucleotide; A, modulator (HKL). **(B)** Binding of carba NAD (c-NAD) to Sirt3.Ac-p53-AMC complex, in presence and absence of 6.25 µM HKL measured using MST.

For this substrate as well, we have verified HKL binding to catalytically active complexes (Fig. 12) and a slight k_cat_ efficiency decrease in the presence of modulator (Table S1). MST measurements on carba-NAD^+^ indicate a decrease in NAD^+^ binding affinity (Fig. 12). In this case, we do not know the catalytic efficiency at higher [NAM] because we have not estimated the *K*_*1*_ parameter. Hence a standard mixed inhibition model was fit instead of Eq. (1).

### Non-steady state activation of SIRT3 by honokiol

The initial rate of deacetylation of the MnSOD substrate was observed to increase in the presence of HKL (Fig. 13). This effect was observed under conditions where the initial rate of deacetylation was sufficiently low that the initial rate could be distinguished from the steady state rate.

**Fig. 13.**
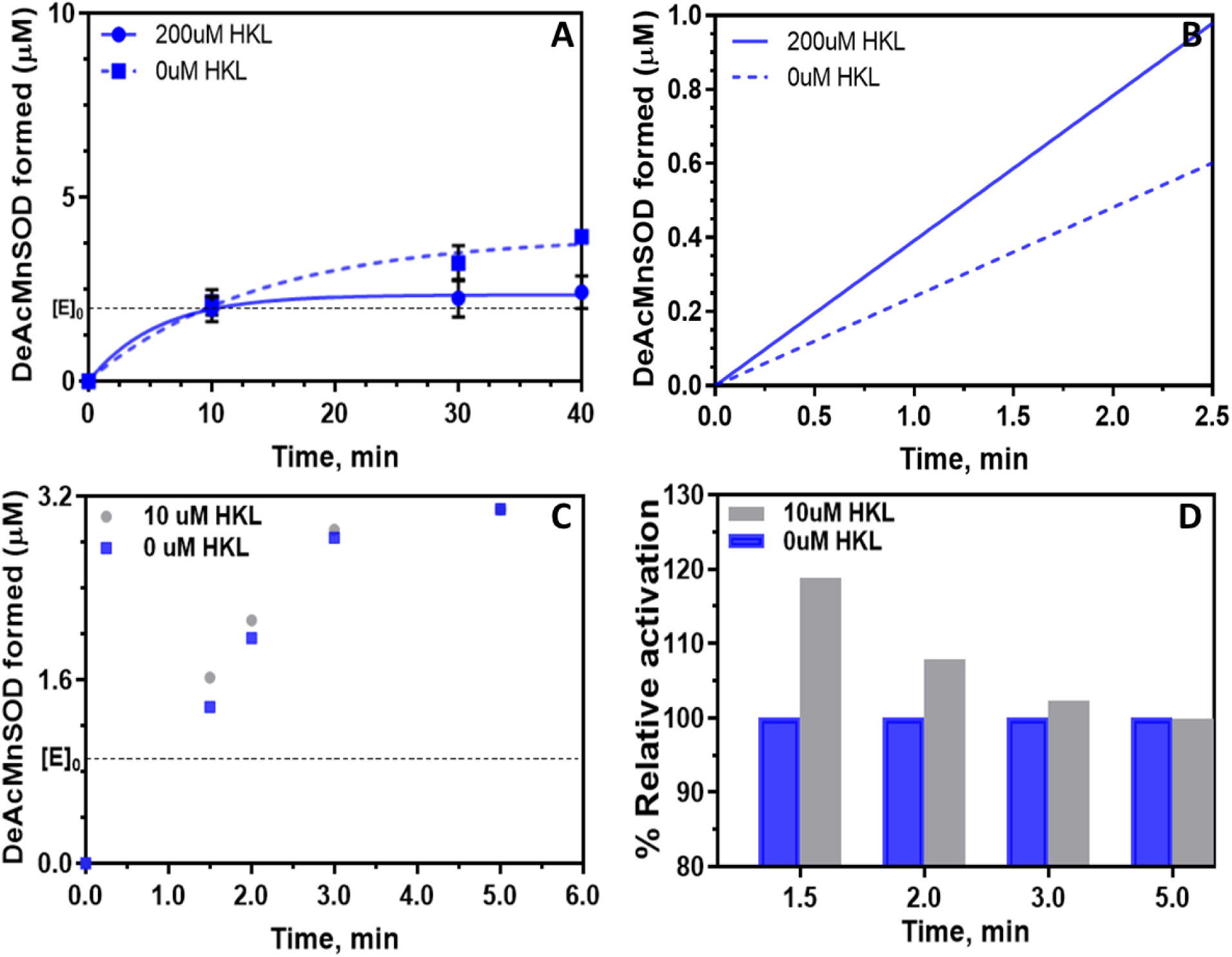
Effect of Honokiol on Sirt3 deacetylation activity: initial rate determination. (**A**) Single exponential fitting of DeAc-MnSOD formed vs. time (0 – 40 min) in the presence and absence of saturating (200μM) [HKL] for 100μM NAD^+^, 100μM NAM; (**B**) Corresponding linear rate fitting plots (initial rates). N=1 for Fig. 13C,D; the error bar (N=6) for t=1.5 min is reported in Fig. 8C. **Sirt3 Activation by HKL. (C)** Plot of product formation vs. time in the presence and absence of 10uM HKL for [NAD^+^]=50uM, [NAM]=0. **(D)** Bar diagram of % relative activity vs. time under the latter conditions. The horizontal dotted line depicts the initial enzyme concentration.

A 0 to 40 minute time series fitting under these conditions captures the initial rate enhancement in the presence of saturating HKL (Fig. 13A,B) whereas a longer time series enabled identification of pre-steady state and steady state phases, and accurate estimation of the steady state rate through double exponential fitting (Fig. 14). The transition between the pre-steady state phase (with HKL-induced activation) and steady state phase occurs after the concentration of deacetylated product exceeds the concentration of enzyme, i.e. after each enzyme molecule has catalyzed one reaction. This pre-steady state effect is also observed in the dose-response curve (Fig. 8A) at 100μM NAD^+^, 100μM NAM ([E]_0_ << [S]_0_ conditions).

**Fig. 14.**
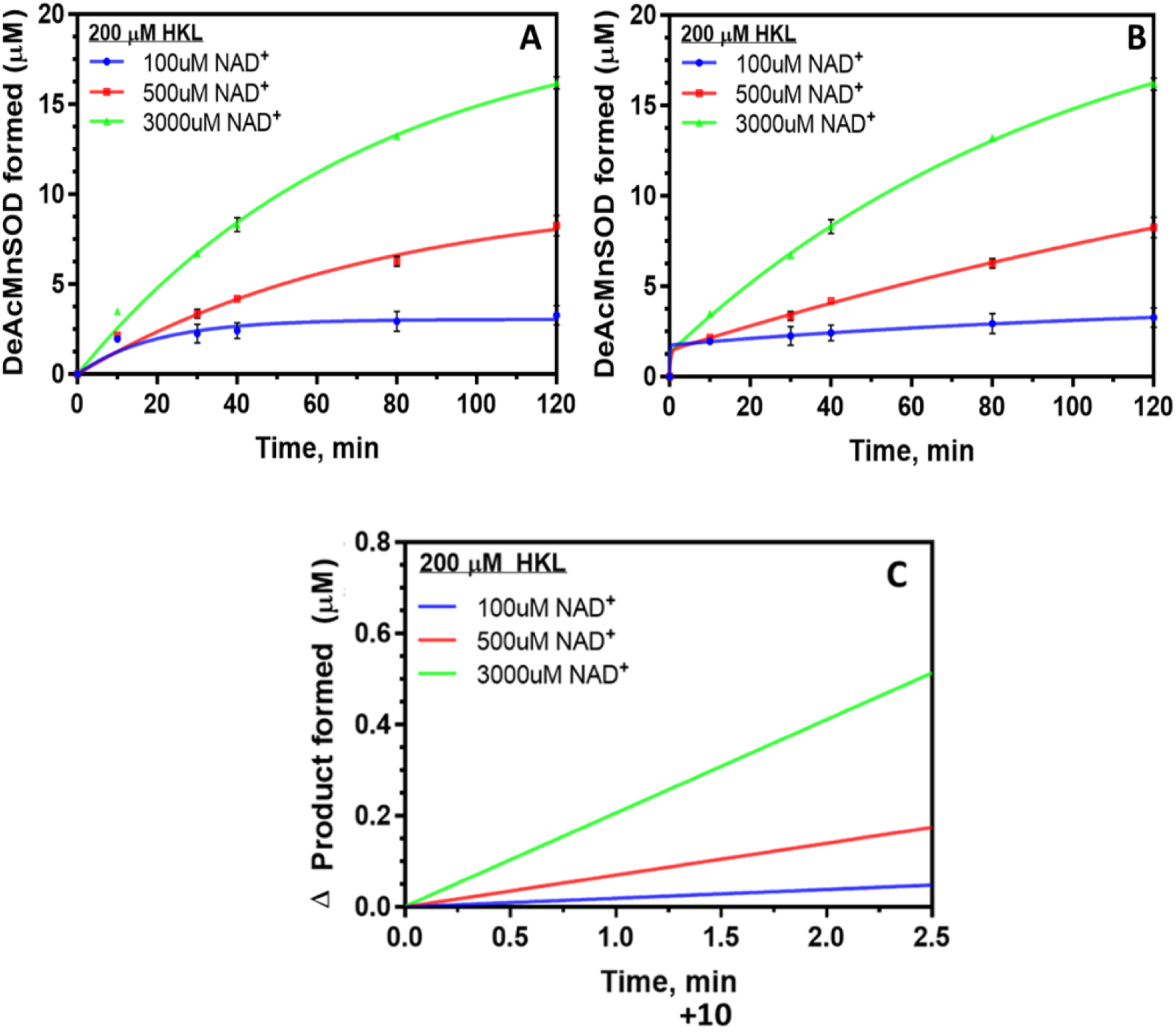
Sirt3 deacetylation activity in the presence of Honokiol for different cofactor (NAD^+^) concentrations: steady state rate determination. Plots of DeAc-MnSOD formed vs. time with **(A)** single exponential fitting, and **(B)** double exponential fitting. Corresponding linear rate fitting plots for **(C)** double exponential (steady state rates) starting at 10 min. [NAM] = 100μM.

Non-steady state activation by HKL was further explored through direct measurements of activity at shorter times (<= 5 min) and also unsaturating HKL. The results are reported in Fig. 13C,D, which depict pre-steady activation of SIRT3-catalyzed deacetylation by HKL through measurements of product formation at short times. While the data in Fig. 13A (and also the dose-response curves in Fig. 8A,B) were collected for the purpose of steady state analysis and characterization, the data in Fig. 13C,D (and Fig. 8C) were collected for pre-steady state analysis. As noted, the use of unsaturating [HKL] (as in Fig. 13C,D) can facilitate activation, but saturating [HKL] (as in Fig. 13A) is required for steady state parameter estimation in the presence of bound HKL. The standard errors of the pre-steady state activation in the presence of 10uM HKL were reported in Fig. 8C above, with p < 0.001.

Thus, honokiol is a non-steady state SIRT3 activator.

### False positive testing of additional hit compounds

Finally, another class of reported sirtuin activators – dihydropyridines, or DHPs (Fig. S7) were previously studied with only labeled assays [21, 23] that may be prone to false positives. We studied the effects of these compounds on SIRT3 activity using the p53-AMC substrate and two assays – both a fluorescence-based and an HPLC assay. The results, presented in the SI (Figs. S8-S10, Tables S2, S3), suggest that DHPs are false positive compounds and not sirtuin activators. The same labeled substrate was used by both Mai et al. [21] and us. In addition, protein-ligand docking did not identify a high affinity binding site. Careful scrutiny should be applied to DHPs reported as activators, due to this finding and the autofluorescence of DHPs (see Fig S10). Ideally, similar protocols for the identification of false positives will be applied to other reported activators for sirtuins other than SIRT1 studied using only labeled assays. Some previous reports of SIRT1 activation for labeled substrates did not properly identify false positives due to the lack of appropriate controls [44]. Labeled assays can be used for high throughput screening as long as appropriate controls and protocols for hit validation are regularly applied.

Application of the rigorous methods for false positive identification and characterization of mechanism-based sirtuin modulators reported herein should facilitate the elimination of false positives in high-throughput screens. The appropriate controls were applied in the current kinetic study of SIRT3-HKL with p53-AMC substrate, which eliminates the possibility of false positives.

### Numerical simulation of activation by MB-STAC hit compounds

As shown above, for the MnSOD substrate, HKL increases the binding affinity of NAD^+^ in the SIRT3 ternary reactants complex, alters the base exchange equilibrium constant, and also increases the binding affinity of the AADPR coproduct. This was shown to increase the pre-steady rate of deacetylation but not the steady state rate under saturating HKL. In order to efficiently analyze the net result of these effects on the dynamics of SIRT3-catalyzed deacetylation over the entire course of the reaction, we carried out numerical simulations of the reaction dynamics under multiple initial conditions including several [E]_0_/[NAD^+^]_0_ at saturating AcPr, over all three phases of the reaction. In these simulations, we applied the simplifying assumptions that a) all ligands have equal off rates in the absence of HKL and NAD^+^ has an on rate 10 times that of NAM; b) HKL increases the on rate of NAD^+^ by a factor of five, increases the rate of ADP ribosylation by a factor of five, and reduces the off rate of AADPR by a factor of 10. The results are shown in Figs. 15 and 16.

**Fig. 15.**
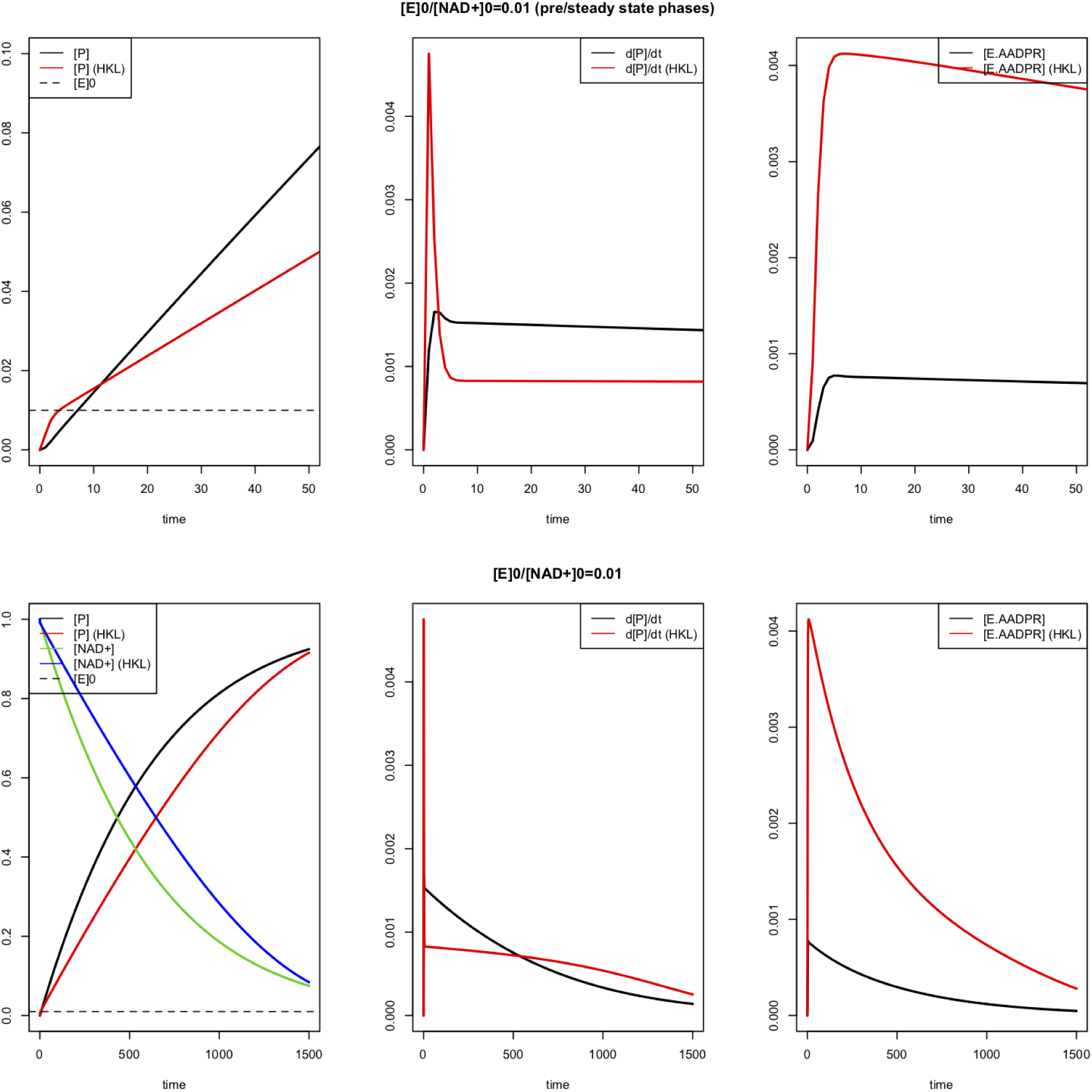
Numerical simulations of the dynamics of SIRT3-catalyzed deacetylation under [E]_0_/[NAD^+^]_0_ << 1 conditions in the presence and absence of a non-steady state MB-STAC like HKL. Top: pre-steady state and steady state phases; Bottom: full reaction. P = deacetylated peptide product. Simulation parameters are defined in the text, and do not precisely correspond to the true parameter values for HKL.

**Fig. 16.**
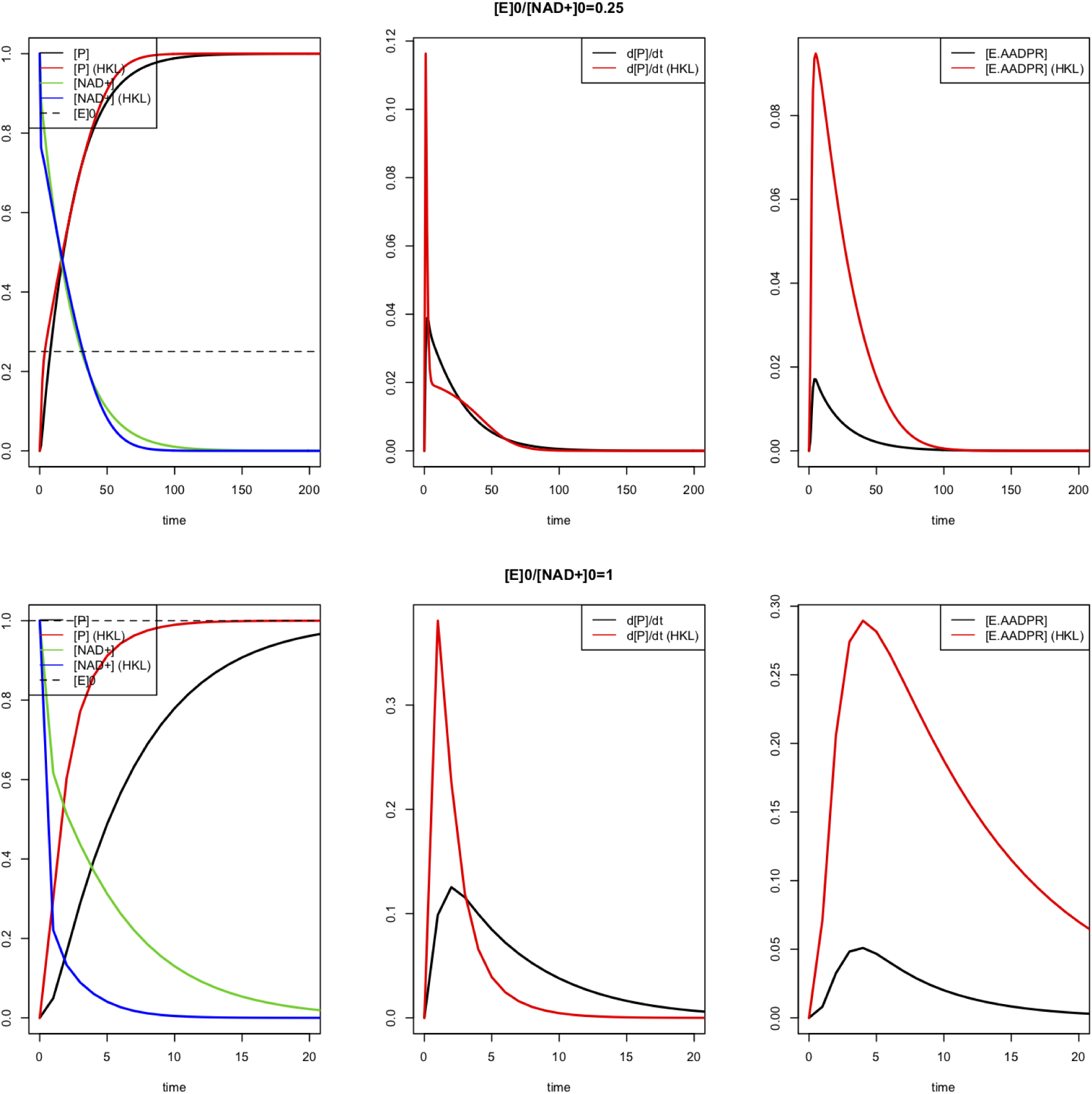
Numerical simulations of the dynamics of SIRT3-catalyzed deacetylation under [E]_0_/[NAD^+^]_0_ < 1 (top), = 1 (bottom) conditions in the presence and absence of a non-steady state MB-STAC like HKL. P = deacetylated peptide product. Simulation parameters are defined in the text, and do not precisely correspond to the true parameter values for HKL.

## DISCUSSION

### Characterization of HKL modulation of SIRT3 using computational, kinetic, thermodynamic data

The above data indicate that HKL cobinds with the peptide substrate and cofactor of SIRT3. Under the conditions tested (saturating HKL), the net effect of SIRT3 on MnSOD peptide substrate was pre-steady state activation followed by steady state inhibition. The mechanistic model above is required to understand the effects of HKL on the kinetic parameters of the enzymatic reaction, explain HKL’s activation of SIRT3 activity under physiological non-steady state conditionsand evaluate the potential to improve HKL’s activation of SIRT3 through hit-to-lead evolution.

First, the computational results in Figs. 4 -7 demonstrate that HKL cobinds with SIRT3 substrates near the active site and induces a change in the conformation of the cofactor binding loop due to the latter’s flexibility. It is not an allosteric modulator. The induced changes in the cofactor binding loop conformation affect the binding of NAD^+^ as well as that of the AADPR coproduct. This conformational change is predicted to have a significantly favorable effect on AADPR coproduct binding affinity (Fig. 7), whereas its effect on NAD^+^ binding affinity is predicted to be less significant. Moreover, due to the interaction of the modulated loop conformation with the nicotinamide moiety of NAM, (Fig. 6), the resulting change in NAD^+^ binding geometry is expected to alter the base exchange rate constants of the reaction.

Next, we consider the effects of HKL on SIRT3 activity against the MnSOD substrate, in the context of traditional enzyme inhibition models. According to Fig. 11B, *K*_*d,NAD+*_ remains roughly unchanged for this substrate, implying that *K*_*d1,A*_ ≈ *K*_*d2,A*_ in Fig. S3. In the traditional picture, HKL binding is noncompetitive with respect to NAD^+^. According to Fig. 11D, the increase in HKL binding affinity in the presence of NAM suggests that *K*_*d,NAM*_ decreases for this substrate, implying that *K*_*d3,A*_ *> K*_*d4,A*_ in Fig. S3. In the traditional picture of enzyme inhibition, HKL binding is uncompetitive with respect to NAM.

However, the effect of HKL on activity can only be understood by application of the full kinetic and thermodynamic model described above, which involves simultaneous effects of the modulator on both the forward and reverse reactions in the context of a steady-state model of the reaction. Note that in the presence of HKL, *K*_*m,NAD+*_ increases and catalytic efficiency decreases by a somewhat larger margin. The parameter estimates in Table 1 together with the full set of MST binding affinity data can be interpreted within the context of the model above to shed light on the mechanism of HKL modulation and to evaluate its properties as a hit compound.

First, note from Table 1 that *K*_*1*_ decreases several fold in the presence of HKL, consistent with the increase in NAM binding affinity suggested by MST measurements. As discussed in [27], a decrease in *K*_*1*_ may be inconsequential in terms of a molecule’s propensity to increase the enzyme’s catalytic efficiency as long as the common assumption of fast NAM release (fast k_-3_ approximation) holds.

Second, *K*_*3*_ decreases several fold in the presence of HKL (Table 1), whereas an increase in *K*_*3*_ is desirable. However, *K*_*1*_*/K*_*3*_ remains roughly unchanged, as can be observed graphically in Figs. 10B, D where the effect of *K*_*1*_ becomes diminished at high [NAM]. This indicates that the reduction in *K*_*d,NAM*_ may be playing an important role in the observed reduction of *K*_*3*._ Under the approximations discussed in [27], the modulated *K*_*ex*_*’* may be relatively close to *K*_*ex*_ (see Fig. S2 for the relationship between *K*_*3*_ and *K*_*ex*_). This is a favorable property of a hit compound since it would imply *K*_*d* 3, *A*_ ≈ *K*_*d* 2, *A*_ ; hit evolution could be carried out to reduce *K*_*d3, A*_, which is conducive to activation. Note that since *K*_*d,NAD+*_ remains roughly unchanged by the modulator, if *K*_*ex*_ is also roughly unchanged, any reduction in k_2_ that affects *K*_*m*_ (see Eq. 2) is associated with a similar reduction in k_-2_.

Third, α decreases several fold in the presence of HKL (Table 1). This shifts the intersection point of the lines in Fig. 9A to the left in Fig. 9B. Under the above hypotheses that *K*_*d* 2, *A*_ ≈ *K*_*d*1, *A*_ *K*_*d* 3, *A*_ ≈ *K*_*d* 2, *A*_, and *K*_*d* 3, *A*_ > *K*_*d* 4, *A*_, this is expected according to the expressions for *K*_*2,app*_ and *K*_*3,app*_ in Appendix A1. Under the approximations described in [27], this decrease in α is due to the fact that *K*_*m*_ increases in the presence of HKL while *K*_*d*_ does not increase (see Eq. 3). We can observe by comparing Figs. 10 C, D to A, B that in the presence of NAM, the extent of inhibition by HKL diminishes, and that due to the reduction in α, this effect is most pronounced at low [NAD^+^], which is the condition under which activation is therapeutically most useful. Under the above hypotheses, through further hit to lead evolution, this property might be exploited to activate the enzyme at low [NAD^+^] in the presence of physiological [NAM] or, going further, to allow a *K*_*m*_ decrease in the absence of product.

Finally, the observed increase in HKL binding affinity in the presence of O-acetylated ADP ribose coproduct (Fig. S4A) may be consistent with the decrease in k_cat_, if product release is rate limiting for this substrate. Also, stabilization of coproduct binding may be consistent with HKL favoring a closed over an open loop conformation; for example, Ex-527, which is known to preferentially bind to a closed loop conformation, reduces coproduct dissociation rate and improves its binding affinity [39].

We can also understand the properties of conventional inhibition plots with respect to modulator concentration based on the mechanistic information gleaned from our studies. Note from Fig. 10A, the intersection point of the lines with and without modulator lies to the left of the y axis and above the x axis. In the traditional mixed inhibition picture of enzyme modulation (which does not distinguish between *K*_*m*_ and *K*_*d*_), this is due to the modulator both increasing *K*_*m*_ and decreasing *v*_*max*_, but this traditional picture does not distinguish between *K*_*m*_ and *K*_*d*_, and also does not account for the relation between *K*_*m*_ and *v*_*max*_ (see Eq. 2 and Fig. S2). By contrast, the equations for the lines in the presence and absence of modulator can be understood in terms of the fundamental dissociation constants and rate constants of the reaction based on our mechanistic analysis above.

Note also from Fig. 10C, the intersection point of the lines with and without modulator moves closer to the x axis in the presence of NAM. For sufficiently high [NAM], the intersection point of these lines falls below the x axis and then will eventually move to the right of the y axis. The fact that HKL does not appear to increase *K*_*d,NAD+*_ gives it higher relative activity at low [NAD^+^] in presence of NAM (this is especially pronounced in the pre-steady state phase, see inset in Fig. 9d). If this could be achieved in the absence of NAM, it would signify an activator that increases catalytic efficiency while reducing *v*_*max*_.

The ability to relate the modulated steady state kinetics of the enzyme to the manner in which a hit compound interacts with the various species in the reaction mechanism is a feature of mechanism-based activator discovery, in contrast to hit validation in traditional drug discovery, where the focus is on binding affinities [10].

Pre-steady state kinetics can provide additional information about the mechanism of modulation, as discussed further below. For enzymes where product dissociation is the rate-limiting step, initial rates may not be similar to steady state rates. Hence, using initial rates in inhibition model fittings can be used for qualitative model selection, not quantitative parameter estimation. In the present work, we applied double exponential time series fits for quantitative work, which accounted for the possibility of a pre-steady state phase affecting initial rates (Fig. 14). Had we fit a single exponential, the estimated *v*_*max*_*s* would be similar but the estimated *K*_*m*_*s* would decrease, increasing the estimated catalytic efficiency in in the presence of HKL beyond what we have reported. Importantly, if slow product dissociation results in the significantly slower turnover rate in the steady state phase, then HKL may not reduce the rate of the slowest deacylation chemistry step in stage 2 of the reaction (discussed further below).

Turning to the p53-AMC substrate, in Table S1, which compares the initial rates of catalysis with this substrate in the presence and absence of saturating HKL, we observe a change in *v*_*max*_ but not *K*_*m*_ -- which would be characterized as noncompetitive inhibition in the traditional picture. However, as discussed above for the MnSOD substrate, the traditional picture is not sufficiently informative and the mixed inhibition plots with respect to NAM provide the important mechanistic information. In the absence of NAM, there is a 3-4 fold decrease in catalytic efficiency for this substrate. However, whereas α decreased in the presence of HKL for the MnSOD substrate, it increases for the p53-AMC substrate. Since for MnSOD substrate, there was no change in *K* _*d,NAD*_ ^*+*^ and a reduction in *α*K*_*m,NAD+*_ induced by HKL, the greater change in *α*K*_*m,NAD+*_ induced by HKL for p53-AMC substrate observed in is consistent with the increase in *K* _*d,NAD*_ ^*+*^ as measured by MST (Fig. 12).

Having identified a hit compound for mechanism-based activation that displays at least some of the qualitative features identified above under the rapid equilibrium segments approximation, we can now use the kinetic and thermodynamic data above -- which together constitute a complete set of measurements -- to simultaneously estimate all the modulated rate constants k_1_’, k_-1_’, k_2_’, k_-2_^’^, k_3_’, k_-3_’, k_4_’ (depicted in the model schematic Fig. S3) and associated free energy changes in the presence of the modulator. The ability to identify system parameters in this manner will allow rapid characterization of mutated hit compounds during hit-to-lead evolution to identify those with a favorable balance of properties suitable for further development [45].

The mechanistic analysis presented herein is necessary to understand effects of hit compounds like HKL on sirtuin activity because the net effect of activation or inhibition depends on precise physiological conditions. The current theory explains this dependence. The dependence on time of the relative rates of MnSOD peptide deacylation with and without HKL (Fig. 13) is relevant to the physiological effects of HKL on SIRT3 deacylation *in vivo*. HKL increases the rate at short times, pointing to the importance of non-steady state effects In particular, the ratio [E]_0_/[S]_0_, where [S]_0_ denotes the initial concentration of the limiting substrate, plays a critical role in determining the relative contributions of non-steady state activating phases and steady state inhibitory phases to the net enzyme activity – with higher values of this ratio increasing the extent of activation. It was observed [22] that SIRT3 expression levels increase in the presence of HKL, which is consistent with our mechanistic analysis as it will offset the effect of product inhibition especially in the limit of low [NAD^+^]. Finally, it is also possible to carry out analogous characterization experiments at unsaturating HKL in the dose response curves (at [E]_0_/[S]_0_<<1), in order to obtain the apparent steady state constants defined in Appendix A1. The use of unsaturating modulator concentrations can allow the modulator to dissociate from the product complex, thus enabling product release. Note that the first-order rapid equilibrium segments approximations above to the steady state constants in the presence of modulator (Appendix A1) account for this and do allow for the possibility of an activity maximum at subsaturating concentrations. These points are all explored in further detail in the sections below.

In summary, HKL binds to all complexes in the sirtuin reaction mechanism, not just the product as in the case of mechanism-based sirtuin inhibitors like Ex-527. The tight binding of HKL to the coproduct does not reduce catalytic efficiency. Compounds like Ex-527 are not hits for mechanism-based activation because they do not bind to any other catalytically relevant complexes and because they reduce product dissociation rate significantly, thus extinguishing the reaction under saturating conditions. Comparing the effects of saturating HKL and saturating Ex-527 [40] on SIRT3 activity, due to the substantial reduction in coproduct dissociation rate (steady state k_cat_ close to 0), saturating Ex-527 effectively constitutes “single hit” conditions wherein each enzyme can only turn over products once. Ex-527 does this for many substrates and sirtuins including AceCS2 and p53-AMC [40], due to the fact that it protrudes into the C pocket and is hence incapable of binding NAD^+^ in a productive conformation. HKL is a hit for both MnSOD and p53-AMC substrates for reasons discussed above.

### Apparent steady state effects of unsaturating concentrations of MB-STAC hit compounds

As described in [27], although saturating [A] is needed to characterize the effects of a MB-STAC hit compound on the chemical and binding rate constants of the reaction, the hit compound can be applied at unsaturating concentration in order to leverage the positive effects of A on certain steps of the reaction while reducing its negative effects on other steps. At unsaturating [A], the reaction can follow a path between the front and back faces of the cube resulting in apparent steady state constants as shown in Fig. S3.

As such, note that the physiological effect of HKL activation was reported at 10μM concentration [22]. At this concentration, according to Fig. 8, there is very little net effect of HKL on activity under steady state conditions. Note that 10μM HKL is above the K_d_ of the compound as displayed in Fig. 11 (also note there is no secondary binding event at higher [HKL] as shown in this graph.) However, the effect of HKL on activity in the steady state dose response curve in Fig. 8 extends above 100μM HKL. This is because the rate of dissociation/association of HKL from/to the enzyme with respect to the other rate constants of the reaction can affect the apparent values of the steady state rate constants in Fig S3 and this effect is essentially eliminated at HKL concentrations above 100μM. On the other hand, unsaturating HKL can allow the apparent steady state and rate constants for steps such as coproduct release to be increased with respect to the values of these constants in the presence of HKL (the rate constants on the back face of Fig. S3). Nonetheless, hit-to-lead evolution and lead optimization of MB-STACs should be carried out at saturating [A] (as applied above) in order to increase the robustness of the MB-STAC with respect to physiological expression level and substrate level differences.

The theory of steady state dose response curves of MB-STAC hit compounds is presented in our recent work [46], which provides dose response equations for unsaturating [A]. We emphasize that the experimental dose response curves above were intentionally collected at saturating HKL since these properties are most relevant to hit-to-lead evolution.

### Non-steady state activation by MB-STAC hit compounds

The steady state dose response curves displayed in Fig. 8 were intentionally measured at low [E]_0_/[NAD^+^]_0_ and at saturating HKL for the purpose of characterization of the effects of HKL on deacetylation reaction parameters. Overall, the reaction goes through 3 phases: i) the first (pre-steady state) phase during which each enzyme molecule catalyzes a limited number of reaction cycles, and which is faster in the presence of saturating HKL (Fig. 13); ii) the second (steady-state) phase during which each enzyme molecule may catalyze many reaction cycles, and which is slower in the presence of saturating HKL; iii) the third (post-steady state) phase which is faster in the presence of saturating HKL, once the concentration of the limiting substrate is similar to or lower than the concentration of enzyme. Depending on the initial ratio of enzyme to limiting substrate in the system (and the ratio of enzyme to HKL, as described in the previous section), the net effect of HKL over the course of the reaction can be either inhibitory or activating, with activation occurring if this ratio is above a threshold value. As the initial ratio of enzyme concentration to limiting substrate concentration increases, the duration of the first two phases decreases and the duration of the third phase increases. If this ratio is not low, steady state rate measurements needed to characterize the effects of HKL are not accurate.

The first two phases have been studied experimentally above. The role of product inhibition in non-steady state activation by HKL is apparent from Fig. 13A,C. It can be seen that activation occurs at times prior to which the amount of product formed is roughly equal to the enzyme concentration [E]_0_.

Fig. 8C, which shows activation with high statistical significance, uses measurements from reaction phase i. The dose response curve for 100uM NAD+, 100uM NAM (Fig. 8A) includes contributions from phases i and ii of the reaction, whereas the other dose response curves have dominant contributions from phase ii (steady state phase). The pre-steady state activation presented in Fig. 13 for the MnSOD substrate (for both saturating and unsaturating HKL) imply that the third phase will also display activation for this substrate even in the absence of HKL dissociation and the ability to follow paths combining the front and back faces of the mechanism-based modulation reaction diagram Fig. S3. Indeed, Pillai et al [22] established the activation of the SIRT3 activity on the MnSOD substrate under larger ratios of enzyme to substrate and enzyme to HKL concentrations.

The results from numerical simulation of SIRT3 reaction dynamics for various values of [E]_0_/[NAD^+^]_0_ (Figs. 15, 16) are consistent with the analysis above of the effects of HKL on the three phases of the reaction, and the pre-steady state activation / steady state inhibition observed experimentally. In Fig. 15 ([E]_0_/[NAD^+^]_0_ << 1), note the pre-steady state activation by the MB-STAC followed by a long steady state phase and finally a post-steady state phase. In Fig. 16 (higher [E]_0_/[NAD^+^]_0_), note the net activation, and also the shorter steady state phases which preclude accurate parameter estimation.

Fig. S12 depicts the reaction diagram corresponding to the front face of Fig. S3 that includes separate representations of the product release and final chemistry (deacetylation) steps for modeling of the reaction under non-steady state conditions. (It can be shown that the addition of such irreversible rate constants does not alter the expressions for the steady state constants applied herein.) In the presence of HKL, k_cat_ in the steady state model does not correspond to the rate determining step under non-steady state conditions; rather, being the rate limiting step of product release and the final chemistry step, it is approximately equal to the rate of product release. Since the rate determining step under non-steady state conditions is the final (deacetylation) chemistry step, the pre-steady state activation (Fig. 13) of SIRT3 by saturating HKL, as reported above, indicates that HKL reduces the rate of product release (by binding tightly to the enzyme.AADPR complex depicted in Fig. S12), but not necessarily the rate of the final chemistry step. Non-steady state (phases i and iii defined above) measurements do not provide information on the effect of a modulator on product release, whereas measurements on the steady state system do not distinguish between the effects of the modulator on the rate of product release vs its effects on the rate of the final chemistry step. Hence the two types of measurements are complementary in the characterization of hit compounds for mechanism-based activation.

Importantly, note that in the expression for the steady state catalytic efficiency k_cat /_ K_m,NAD+_, which is relevant to steady state sirtuin activation under NAD^+^ depletion conditions associated with aging, the rate constants in k_cat_ do not explicitly appear. Thus the reduction in k_cat_ by such a modulator does not affect catalytic efficiency. Hence the non-steady state activation by HKL is relevant to the goal of increasing catalytic efficiency, but further hit-to-lead evolution is required.

Note that in order to characterize the mode of action of the mechanism-based inhibitor Ex-527 [40], standard inhibition models were not sufficient, requiring reference to crystal structures. We have presented quantitative methods for the characterization of mechanism-based activator hits that are suitable for high-throughput studies. These data can be applied to the system identification of the effects of HKL on all rate parameters, through a combination of the steady state analysis reported above and time series analysis on the non-steady state kinetic data, as described in reference [45].

## CONCLUSIONS

We have applied a biophysical framework for activation of sirtuin enzymes to the characterization of several proposed nonallosteric sirtuin activators of the SIRT3 enzyme. We presented and analyzed results from computational, kinetic and thermodynamic studies on these modulators. Two different peptide substrates were studied to show how the substrate dependence of mechanism-based activation can be explored using the present framework.

We have shown that the compound honokiol, previously reported to be a SIRT3 activator [22], does activate the enzyme under physiologically relevant non-steady state conditions but does not activate it under steady state conditions. By applying a) computational modeling; b) experimental non-steady state, steady state and thermodynamic assays, and using the theory of mechanism-based enzyme activation to interpret the data, we explained the mechanism behind honokiol’s enhancement of SIRT3’s non-steady state deacetylation rate, demonstrated it is a nonallosteric modulator, and established a foundation for further hit-to-lead evolution of honokiol and related compounds in order to convert them into more potent, steady state activators. As such, the present study constitutes an important foundation for the development of a new class of drugs for the treatment of age-related diseases that operate through a wholly new mode of action not shared by any existing drug.

The enzyme activation theory that was applied herein to these modulators motivates computational and experimental workflows for the hit identification, hit-to-lead evolution, and lead optimization of mechanism-based activators. In particular, the theory enables the identification of important hits that activate enzymes in a dose-dependent fashion or under non-steady state conditions, and evolution of these hits into more robust steady state activators. This is achieved by separating the observed kinetic and thermodynamic effects of a modulator into components and identifying those properties that require optimization through hit-to-lead evolution.. Hit-to-lead evolution could be applied, for example, to HKL modulation of the deacylation rate of specified SIRT3 substrates. Such workflows would bear more similarity to the directed evolution of enzymes than to traditional drug discovery workflows. Future work can apply enzyme engineering-inspired methods to hit-to-lead evolution of nonallosteric activators, in conjunction with structure-based computational methods analogous to those developed for computational enzyme design [47].

## AUTHOR CONTRIBUTIONS

RC designed the experiments; XG, AU and SM did the experiments and analyzed data. RC, AU and XG wrote the manuscript.

## Appendix A1

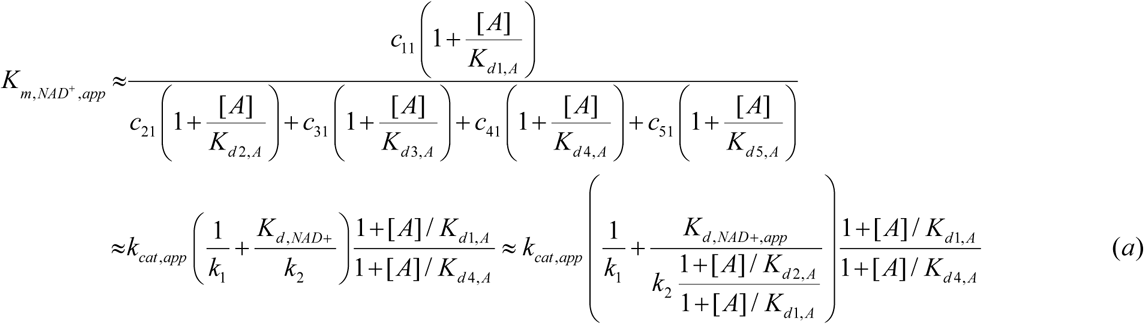

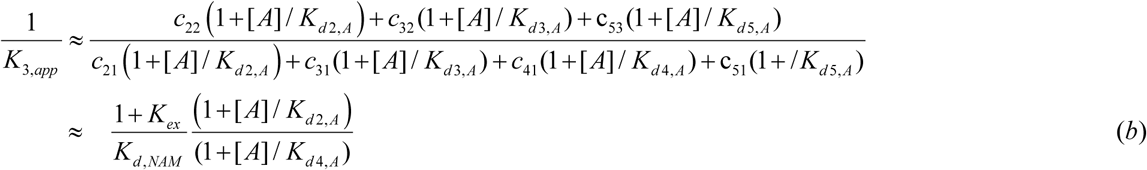

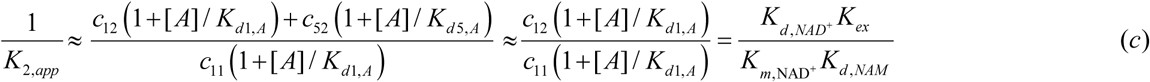

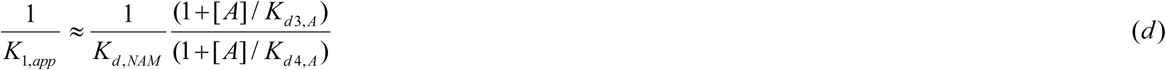

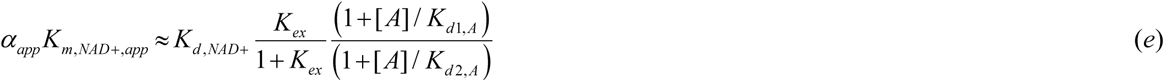

## Appendix A2

Expressions for c_ij_’s in Appendix A1 and Eq. (S1):

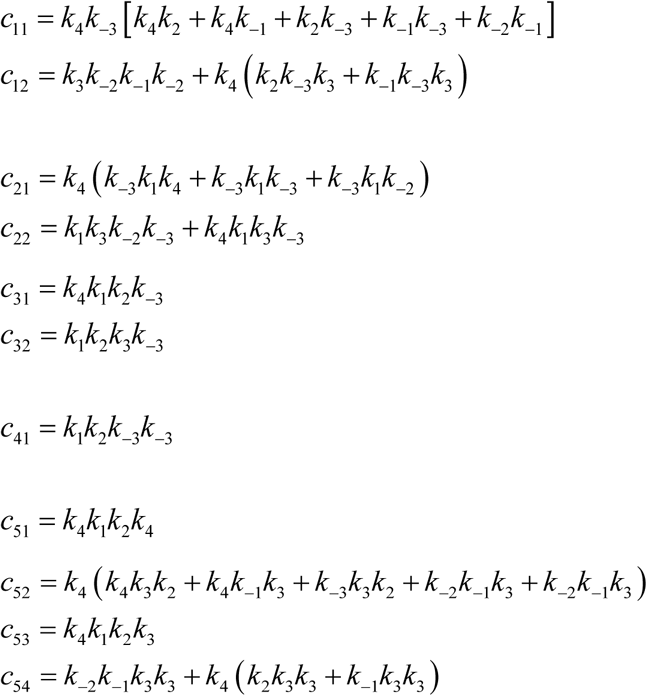

## Supporting Information

### Side chain prediction

Rotamer-based side chain prediction was applied following loop replacement. A subset of residues was chosen for side chain prediction. The residues considered for prediction are 144-180, 195, 199, 204, 207, 210, 227-234, 248, 251, 291, 294, 324 -- residues with 7.5 Å of a modelled loop region were refined. Native side chain conformations were repredicted in native environment prior to prediction in non-native environments.

Four methods for side chain prediction in the protein local optimization program (PLOP) were used:

- Default method -- No backbone sampling or reorientation of the CA-CB bond is performed.
- Monte Carlo approach -- Monte-Carlo sampling of side-chain conformations including backbone flexibility.
- CA-CB vector sampling -- varying the orientation of the CA-CB bond by up to 30 degrees from the initial direction.
- Backbone sampling -- Sample the backbone on a set of 3 residues centered on the residue for which the side chain is being refined.

We used the top 3 conformations provided by the Monte Carlo run for minimization.

### Molecular dynamics

#### Equilibration vs production simulations

There were three phases in the molecular dynamics simulations. First, in the relaxation phase, the system underwent a 2000-step minimization before a short 200 ps NPT molecular dynamics simulation, with the main chain atoms of the protein restrained to the positions in the crystal structures with force constants of 5 kcal mol^-1^ Å^-2^. Next, the systems ran for various lengths of time up to 22 ns in the equilibration phase. Last, the sampling phase included a 10 ns of molecular dynamics simulation. molecular dynamics (first stage): Before launching production run we minimize and equilibrate (200 ps of equilibration) the system and check for the stability (time vs RMSD, time vs potential energy, time vs pressure, time vs temp) to ensure that the system is stable before launching the production run (15 ns). This entails a slow heating up (from 15 k to 300 K with a gradual increase of 15 K) of the system and then running equilibration simulation for 200 ps at 300K.

### Estimation of conformational and binding energies

Energies were calculated using the MM-PBSA and the MM-GBSA methods as implemented in the AMBER package [1]. In MM-PBSA and MM-GBSA, binding free energy is evaluated as:

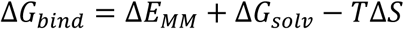

where Δ*E*_*MM*_, Δ*G*_*solv*_, *and T*Δ*S* are the changes of gas-phase interaction energy, solvation free energy, and solute conformational entropy change upon binding, respectively. Δ*E*_*MM*_ includes internal energy in bonded terms, electrostatic and van der Waals energies. Δ*G*_*solv*_ is the sum of polar contributions calculated using the PB or GB model, and nonpolar contributions estimated from solvent-accessible surface area (SASA).

All the calculations were carried out using the MMPBSA.py module with AmberTools13 [1]. The polar contribution of the solvation free energy was calculated by the GB model developed by Onufriev et al. [2] and by the PB method implemented in the pbsa program. A salt concentration of 0.1 M was used in MM-GBSA calculations. The solvent-accessible surface area was evaluated using the LCPO method [3]. Because relative free energy trends were of interest, solute conformational entropy change was neglected.

Energies were evaluated using 10000 snapshots extracted from the last 10 ns at a time interval of 1 ps for each trajectory after ensuring that each one of these trajectories was completely stable.

### Preparation of SIRT3.Ac-Pr.NAD^+^ substrate complex for protein-ligand docking

#### MD simulation protocol

The SIRT3-ALY-NAD^+^ complex is subjected to molecular dynamics simulation. Initially, the complex is kept into a suitably sized box, of which the minimal distance from the protein to the box wall was set to 10A. The box is solvated with the SPC (simple point charge) water model. The counter ions are added to the system to provide a neutral simulation system. The whole system was subsequently minimized by using OPLS2005 Force field. The charges of the atoms of NAD^+^ and ALY were calculated by using OPLS2005 Force field. Covalent and non-bonded parameters for the ligands atoms were assigned, by analogy or through interpolation, from those already present in the OPLS2005 force field. MD simulation is carried out using the Desmond package (version 2.3) with constant temperature and pressure (NPT) and periodic boundary conditions. The OPLS2005 force field was applied for the proteins. The default Desmond minimization and equilibration procedure was followed. Simulations were kept at constant pressure (1 bar) and tem perature (300 K) maintained with a Berendsen barostat and thermostat, respectively. SHAKE was applied to all systems allowing a 2-fs time-step. Long-range interactions were treated with the Particle Mesh Ewald method for periodic boundaries using a nonbonded cut-off of 9.0 and the nonbonded list was updated frequently using the default settings. Coordinates and energies for the OPLS2005 force field simulations were total of 15ns. Extract the structure (15ns) of the equilibrium state from the MD. Het molecules except for native ligands were eliminated (water and counter ions for MD). Minimization is performed to use the OPLS3e force field, which specifies this structure as the reactant structure.

### Separation of MnSOD peptide

An Agilent 1260 infinity HPLC system and a ZORBAX C18 (4.6×250 mm) column were used to separate deacetylated peptide and acetylated substrate using gradient comprising 10% acetonitrile in water with 0.005% TFA (solvent A) and acetonitrile containing 0.02% TFA (solvent B) 1 ml/min flow rate.

### Separation of FdL2 (p53-AMC) peptide

Same column was used with Beckman System Gold HPLC to separate deacetylated product and substrate using gradient comprising 0.05% aqueous TFA (solvent A) and acetonitrile containing 0.02% TFA (solvent B) using 1ml/min flow rate.

In both the cases, upon sample injection, the HPLC was run isocratically in solvent A for 1 min followed by a linear gradient of 0-51 %B over 20 min. After separation, the columns were washed with 100% solvent B and then equilibrated with 100% solvent A. The detector was set at 214 nm. The percent product produced was calculated by dividing the product peak area over the total area. GraphPad Prism software was used (GraphPad Software, Inc, CA) to fit the data.

### Solubility Measurements

Solubility of DHP-1, DHP-2, and Honokiol were measured using HPLC (Agilent 1100 series). In brief, calibration curves were established, for each compound, using concentration range covering the estimated solubility’s. The samples were prepared by adding known amount of the compounds in HDAC buffer containing a range of DMSO. The samples were allowed to equilibrate at 25^0^C for 48 hours before analyzing on calibrated HPLC. Over-saturated samples were prepared by adding excess compounds into the solvent mixtures of interest. The linearity was measured by R-values at least >0.99 and the estimated detection limit was around 0.002 mg/mL (2 μg/mL) based on acceptable N/S ratio.

### Carba-NAD synthesis

Carba-NAD [1((1R,2S,3R,4R)-4-((((((((2R,3S,4R,5R)-5-(6-Amino-9H-purin-6-yl)-3,4 dihydroxytetrahydrofuran-2-yl) methody)hydroxphosphoryl)oxy)oxidophosphoryl)oxy)methyl)-2,3-dihydroxycyclopentyl)-3-carbamodylpryridin-1-ium.] was synthesized according to the method described previously [4].

Briefly, the commercially available (1R, 4S)-2-azabicyclo[2.2.1]hept-5-en-3-one (R(-)-Vince lactam), was asymmetrically dihydroxylated to provide the corresponding dihydroxy lactam after removal of the all-cis stereoisomer. Acid catalyzed methanolysis of the lactam was then carried out to give the amino ester hydrochloride. This was followed by ester reduction using lithium triethylborohydride for reduction of the ester to the alcohol. Next, the nicotinamide carba-riboside was prepared from the alcohol and 3-carbamoyl-1-(2, 4-dinitrophenyl)pyridine-1-ium chloride using sodium acetate in methanol (Zincke reaction). Phosphorylation of the nicotinamide carba-riboside with POCl_3_ according to Lee’s procedure provided the carba-nicotinamide mononucleotide [5]. This was coupled with adenosine 5’-monophospho-morpholidate using pyridinium tosylate and MnCl_2_ as a divalent cation source to create the final pyrophosphate bond.

### General method for DHP-2 synthesis

The synthesis of DHP-2 was carried out by a method described previously [6]. Briefly, in step 1, we first synthesized the core compound, 3, 5-dicarbethoxy-4-phenyl-1, 4-dihydropyridines, by cyclocondensation between benzaldehyde (9.81 µM), ethyl propiolate (9.92 µM) and the cyclopropylamine (14.49 µM). All three compounds were mixed together and heated at 80°C for 30 minutes in 0.5 ml glacial acetic acid. The reaction mix was allowed to cool to room temperature then mixed with 20 ml ddH_2_O for 60 minutes. The resulting solid product was filtered and washed with ethyl alcohol (DHP-1). To obtain 3, 5-dicarboxy derivatives of DHP-1, we performed alkaline hydrolysis of the compound overnight at 80°C in presence of 100% ethanol. After the completion of hydrolysis, the solvent was evaporated, the residue was eluted with water (30 mL), and the resulting solution was acidified with 2N HCl. The precipitate was filtered, washed with 30 ml ddH_2_O three times then dried to obtain pure DHP-2. The compound was then recrystallized by acetonitrile/methanol. The purity and integrity of the intermediate and final compounds were assessed by NMR.

### Effect of DHP-2 on hSirt3^102-399^ deacetylation activity using fluorolabeled peptide

To assess the effect of DHP-2 on hSirt3^102-399^ deacetylase activity, we incubated 5U hSirt3 (0.194 µM) with 3 mM NAD^+^ and 3 µM p53 derived fluorolabeled peptide (FdL2 peptide, Enzo Life Sciences Farmingdale, NY, USA) or 10 µM NAD^+^ and 250 µM peptide substrates or 500 µM NAD^+^ and 250 µM peptide substrates with varying concentrations of DHP-2 (0-400 µM) in a buffer containing 50 mM Tris-HCl, pH 8.0, 137 mM NaCl, 2.7 mM KCl, 1 mM MgCl_2_, and 1 mg/mL BSA. The reactions were carried out for 30 minutes at 37°C and terminated by adding 1X Developer (Enzo Life Sciences). The fluorescence produced was measured (Excitation/Emission: 355/455 nm) using TECAN plate reader (TECAN Infinite M200 PRO, Switzerland, Tecan Group Ltd).

### Effect of DHP-2 on hSirt3^102-399^ deacetylation activity by HPLC assay

Sirt3 enzyme reactions were performed to assess the effect of DHP-2 using p53 derived peptide. The reactions were performed with 3 mM NAD^+^ and 3 µM p53 derived unlabeled peptide or 10 µM NAD^+^ and 250 µM peptide substrates or 500 µM NAD^+^ and 250 µM peptide substrates in the presence of different concentrations of DHP-2, ranging from 0 - 400. The reaction buffer contained 50 mM Tris-HCl, 137 mM NaCl, 2.7 mM KCl, and 1 mM MgCl_2_, pH 8.0. The reactions were started by addition of 5U hSirt3^102-399^ and incubated at 37°C for 30 minutes. The reactions were terminated by adding 2% TFA final then were resolved on C18 column as described above.

**Fig. S1.**
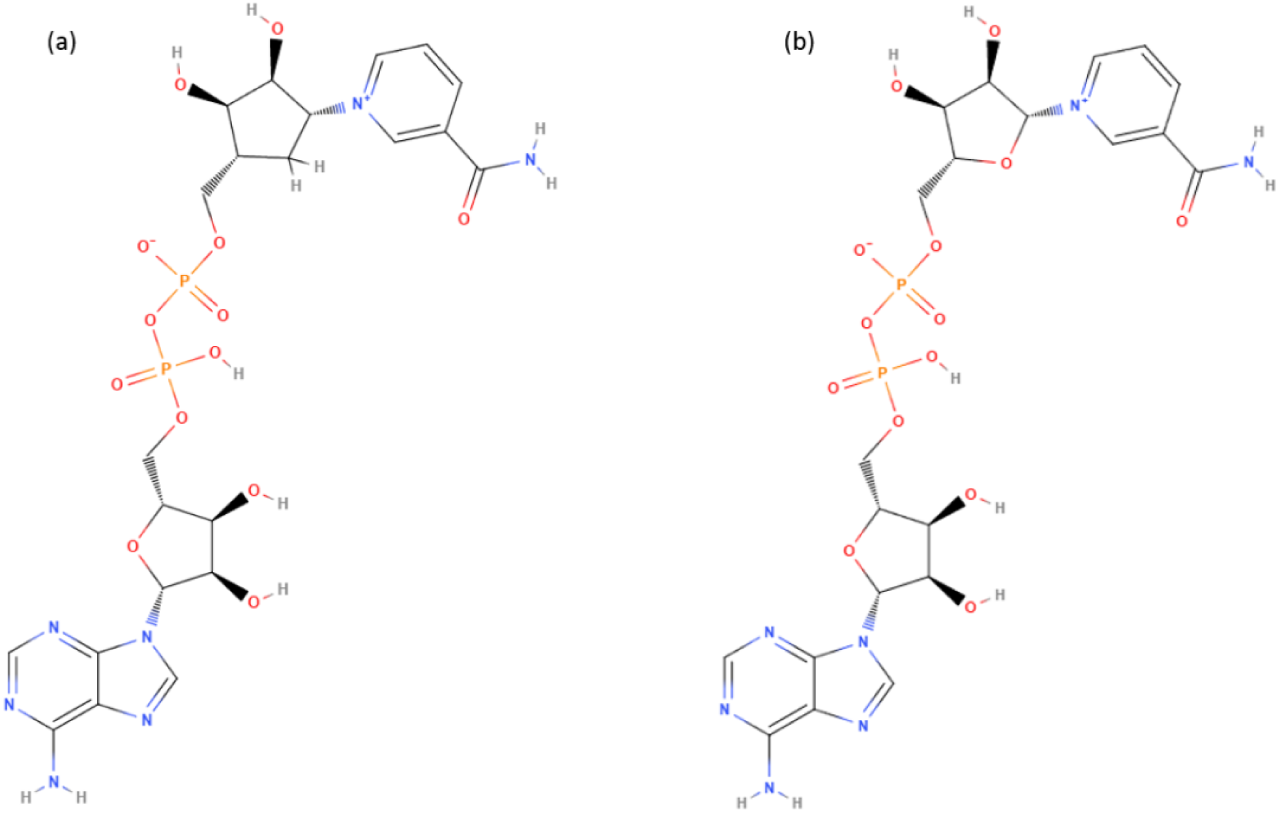
Nicotinamide adenine dinucleotide chemical structures. (**A**) c-NAD^+^, (**B**) NAD^+^

### Steady state constant and rapid equilibrium segments expressions

The rate equations for the reaction network for sirtuin deacylation enable the derivation of steady-state conditions for the reaction. Solving the linear system of algebraic steady-state equations and mass balance constraints for the concentrations [E.Ac Pr], [*E*.Ac Pr.NAD], [*E*.*ADPR Ac* Im.NAM], [*E*.ADPR *Ac* Im], [*E*.*NAM*] in terms of the rate constants and [NAD^+^], [NAM], which are assumed to be in significant excess and hence approximately equal to their initial concentrations [NAD^+^]_0_, [NAM]_0_ respectively, we obtain expressions of the form (A):

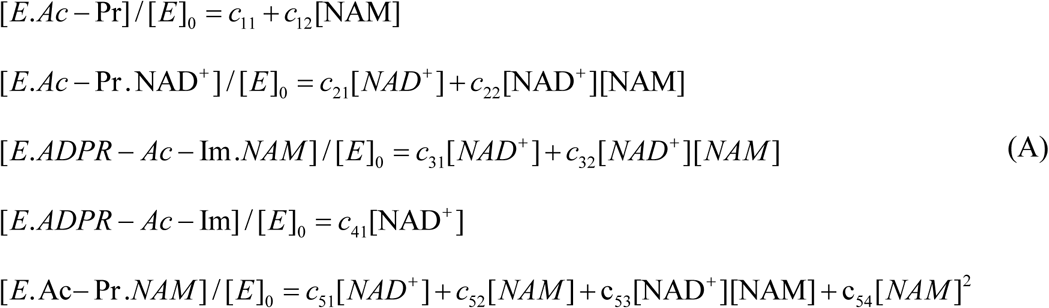

where the term c_54_ that is second order in [NAM] will be omitted from the analysis below. Expressions for the c_ij_’s are provided in the Appendix A2.

**Fig. S2.**
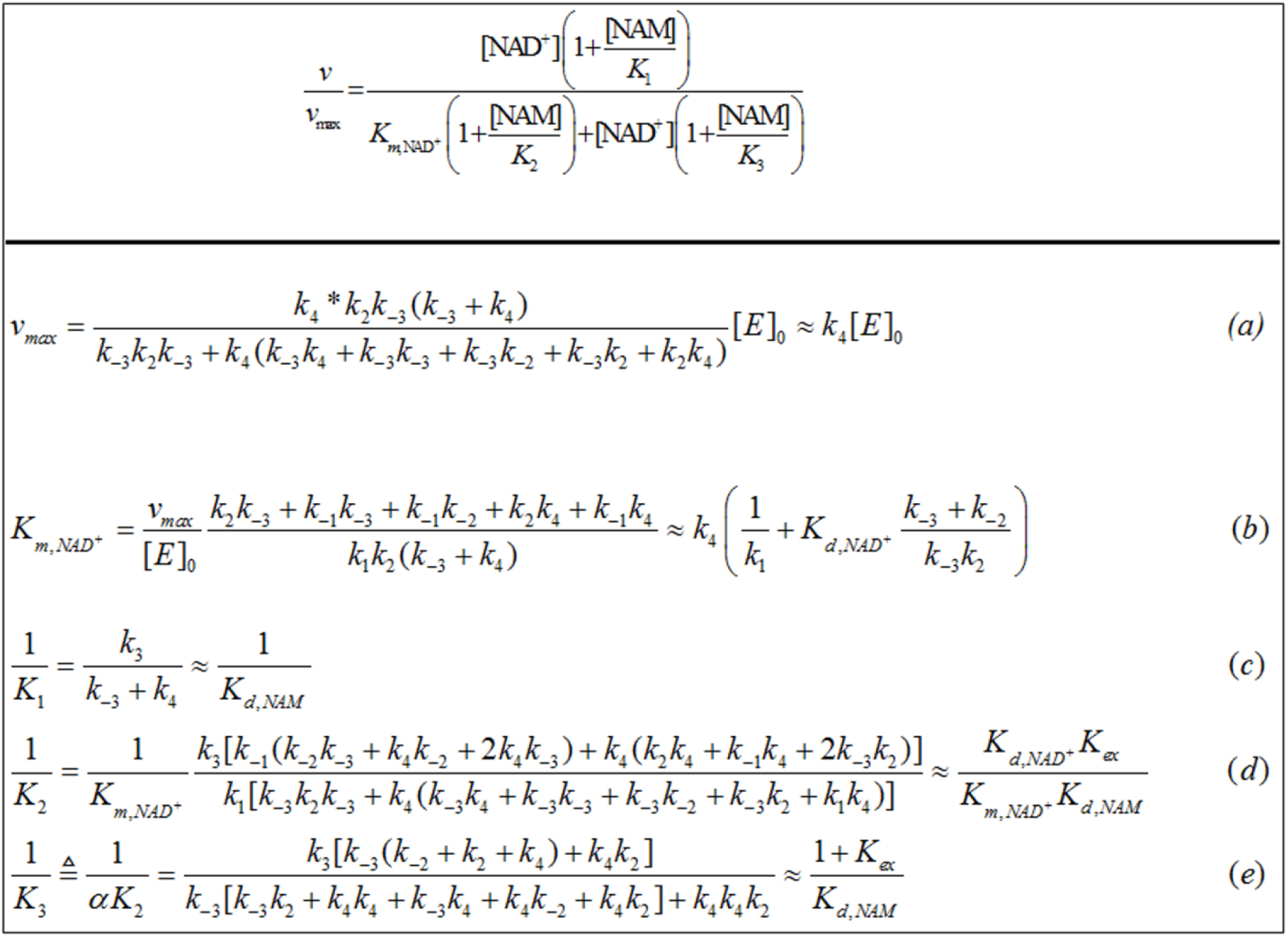
Steady state parameters of sirtuin-catalyzed deacylation expressed in terms of rate constants in the enzymatic reaction mechanism. *K*_*ex*_ = k_-2_/k_2_, and the approximations refer to the case where *k*_*4*_≪*k*_*j*_, *j*≠ 4. Please note that the quality of this approximation can be assessed for the chemistry steps based on QM/MM simulation data, which was cited in the text for yeast and bacterial sirtuins, or from experimental methods described in the text for the estimation of all rate constants in the model.

The expressions for the steady state constants in Fig. S2 (a-e) follow from

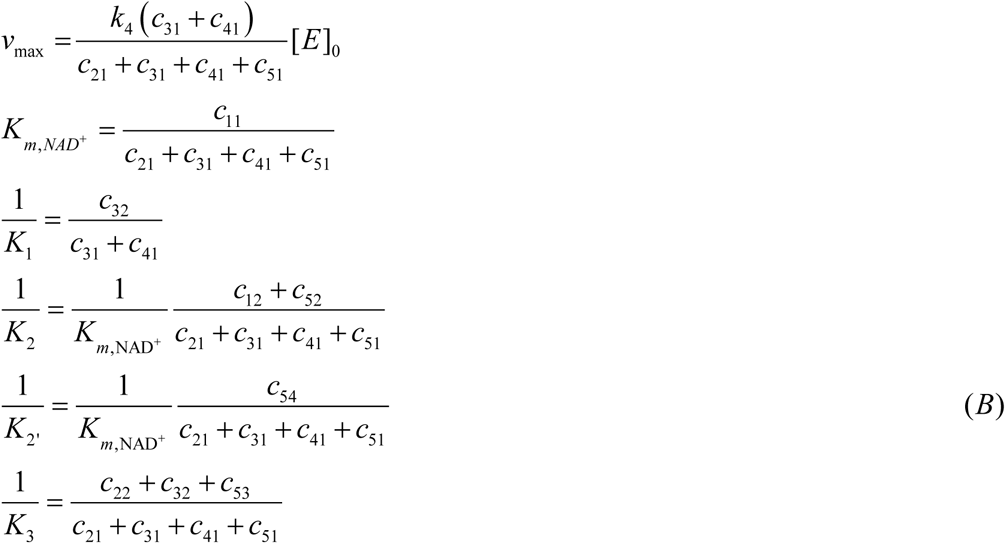

The rapid equilibrium segments expressions for the steady-state concentrations of the species in the sirtuin reaction mechanism depicted in Fig C are:

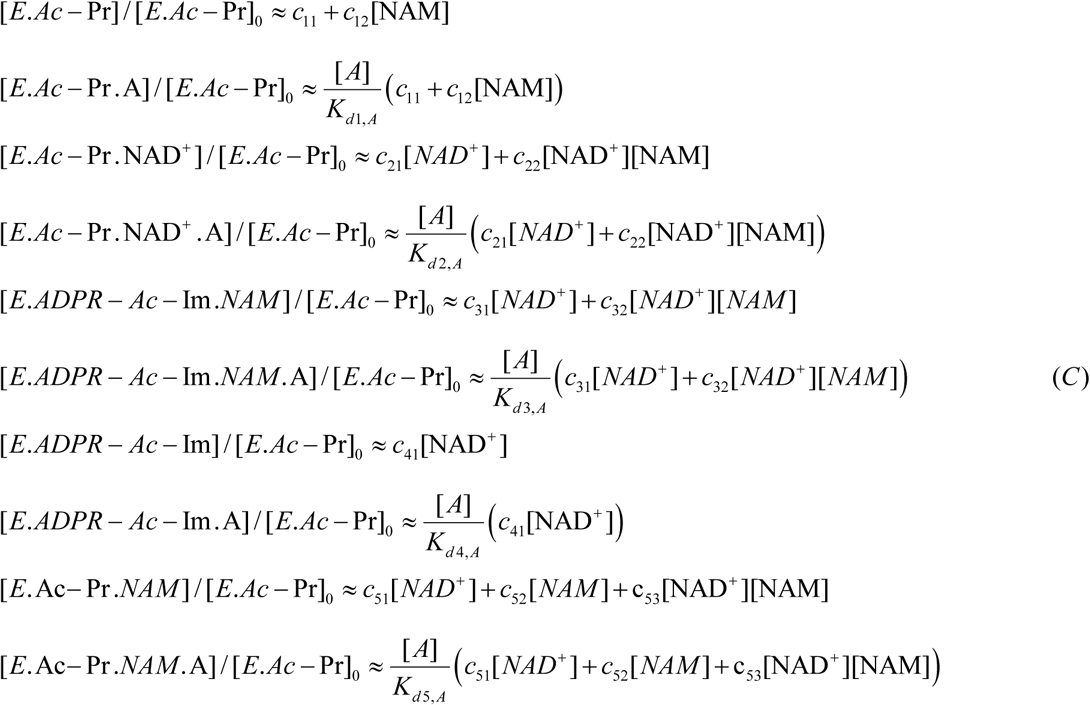

**Fig. S3.**
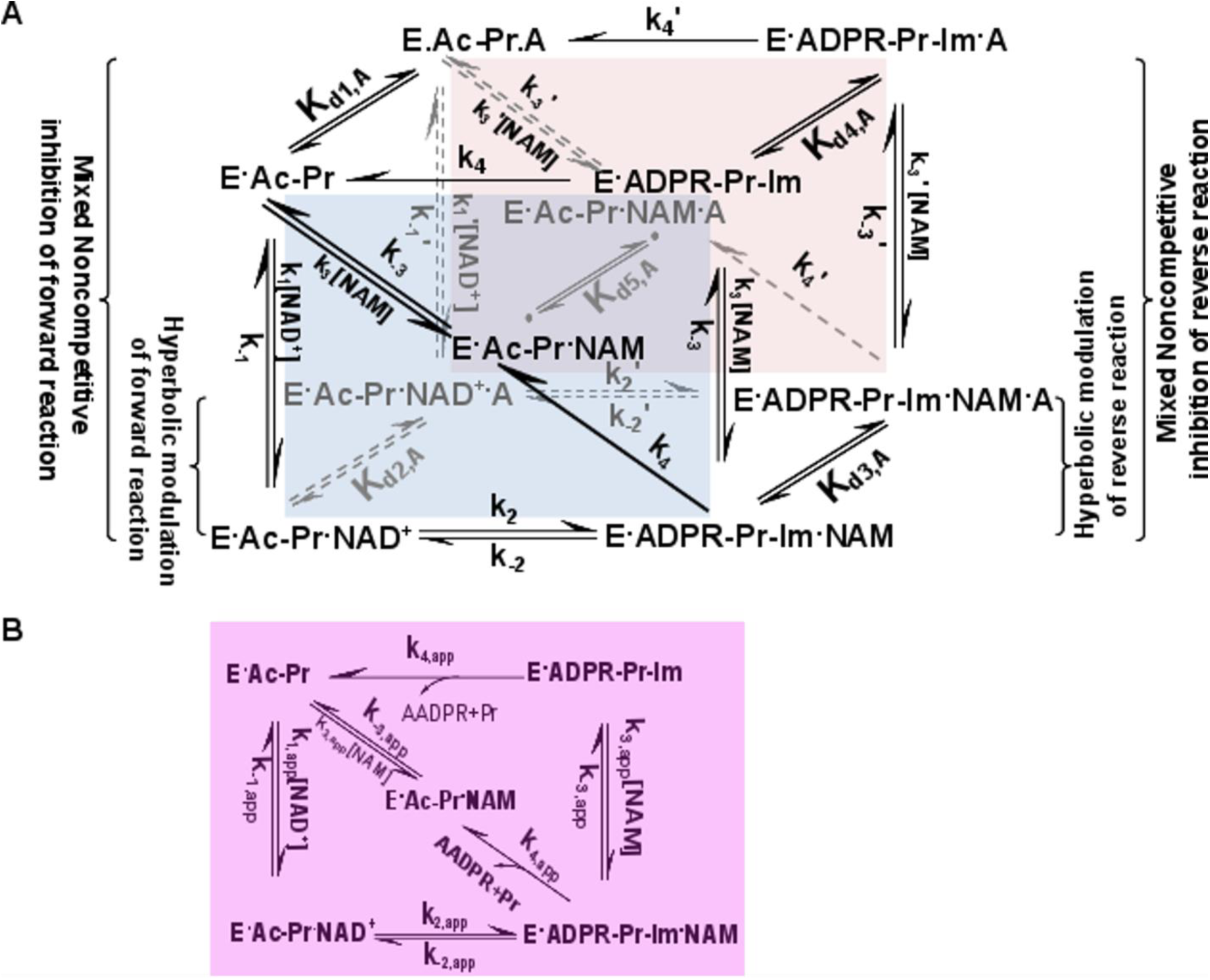
General model for mechanism-based sirtuin enzyme activation. **(A)** The front face of the cube (blue) depicts the salient steps of the sirtuin reaction network in the absence of bound modulator. The back face of the cube (red) depicts the reaction network in the presence of bound modulator (denoted by “A”). Each rate constant depicted on the front face has an associated modulated value on the back face, designated with a prime, which is a consequence of modulator binding. **(B)** The purple face is the apparent reaction network in the presence of a nonsaturating concentration of modulator.

**Fig. S4.**
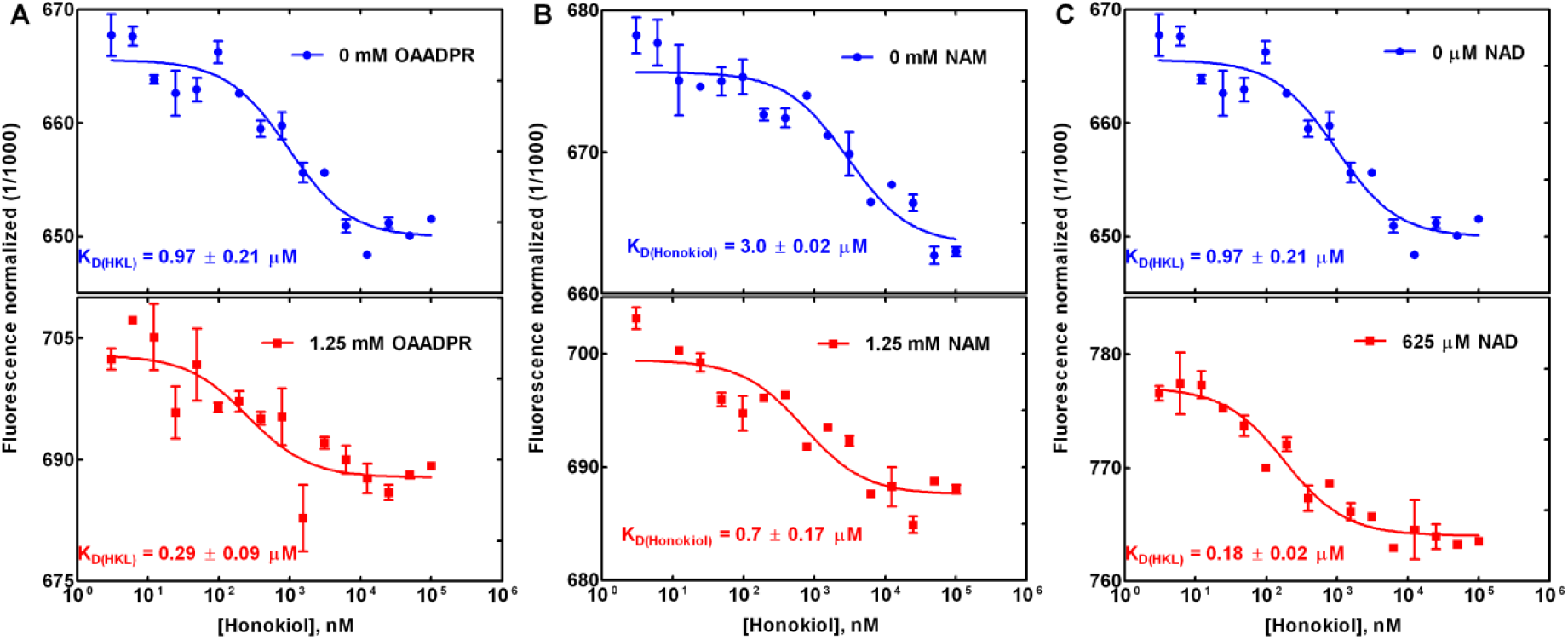
Binding affinity measurements of **(A)** Honokiol binding in the product complex: effect of 2’-O-acetylated ADP ribose coproduct; **(B)** Honokiol binding in the coproduct complex: effect of NAM; **(C)** Honokiol binding in the product complex: effect of NAD^+^

### Substrate dependence of HKL modulation

**Table A.** Model parameter estimates from global nonlinear fitting of mixed inhibition for SIRT3-FdL2 in the presence and absence of HKL. [E_0_] = 3.05 µM

| | 0 $\mu\text{M}$ HKL | 200 $\mu\text{M}$ HKL |
| --- | --- | --- |
| Vmax | 0.3923 | 0.08347 |
| Alpha | 4.007 | $\sim 3.727\text{e}+107$ |
| Ki | 24.52 | 18.14 |
| Km | 1214 | 846 |
| **Std. Error** |  |  |
| Vmax | 0.01506 | 0.002208 |
| Alpha | 2.111 | ~ |
| Ki | 4.154 | 3.388 |
| Km | 107.5 | 61.85 |
| **95% CI (profile likelihood)** |  |  |
| Vmax | 0.3597 to 0.4311 | ??? |
| Alpha | 1.373 to 32.9 | (Very wide) |
| Ki | 17.05 to 38.51 | ??? |
| Km | 986.7 to 1500 | ??? |
| **Goodness of Fit** |  |  |
| Degrees of Freedom | 8 | 8 |
| R square | 0.9961 | 0.996 |
| Absolute Sum of Squares | 0.0002103 | 0.00001222 |
| Sy.x | 0.005127 | 0.001236 |

**Fig. S5.**
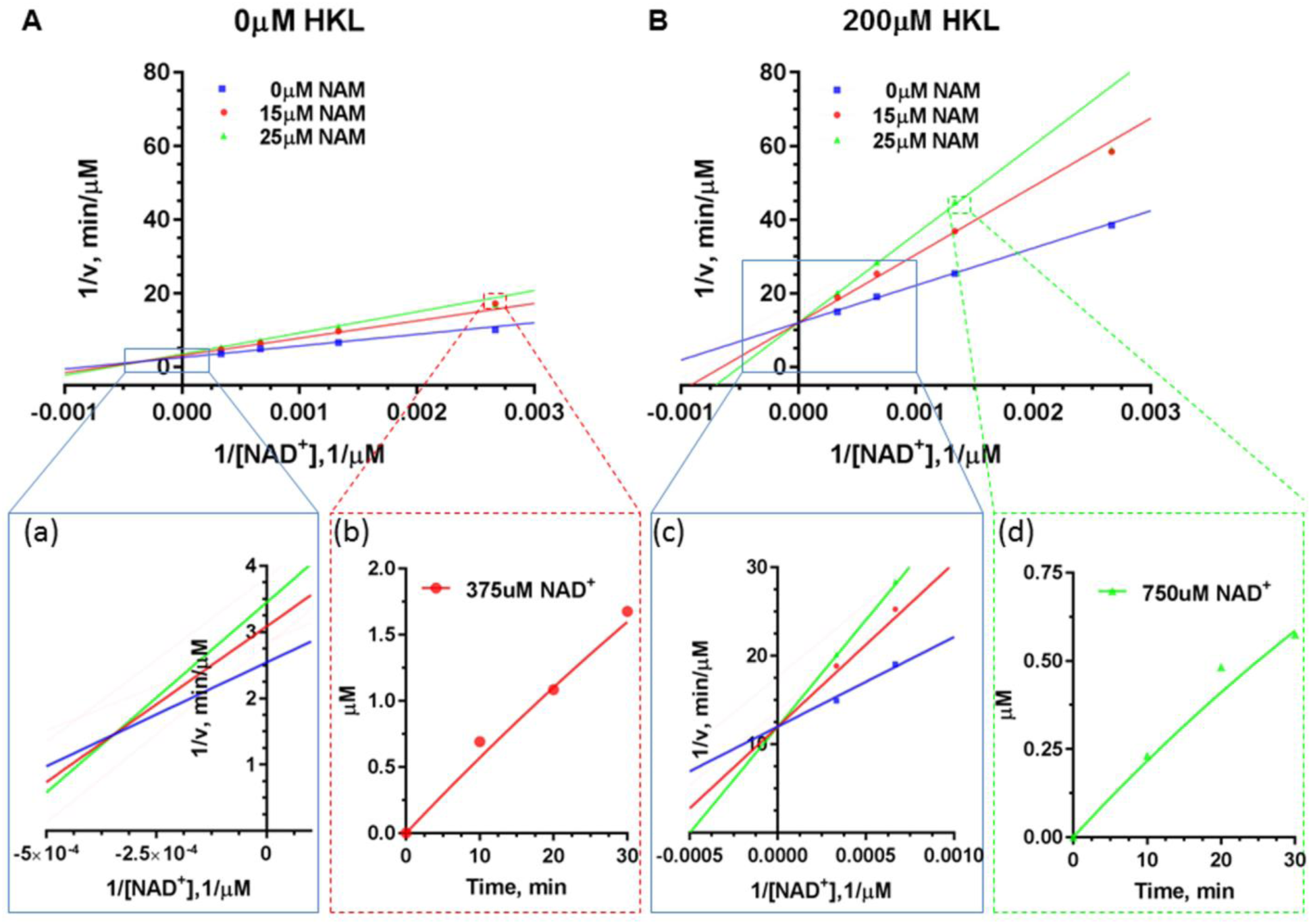
Double reciprocal plots for deacylation initial rate measurements of saturating substrate peptide (FdL2) in the presence of different concentrations of NAM with (A) 0 μM HKL; (B) 200 µM HKL. The enlargement of the intersection points were provided as insets (a) and (c). The time series plot of mM product formed vs. time are provided as insets (b) and (d). [Originally presented in R. Chakrabarti, Mechanism-based enzyme activating compounds, J. Theor. Bio (submitted)]

**Fig. S6.**
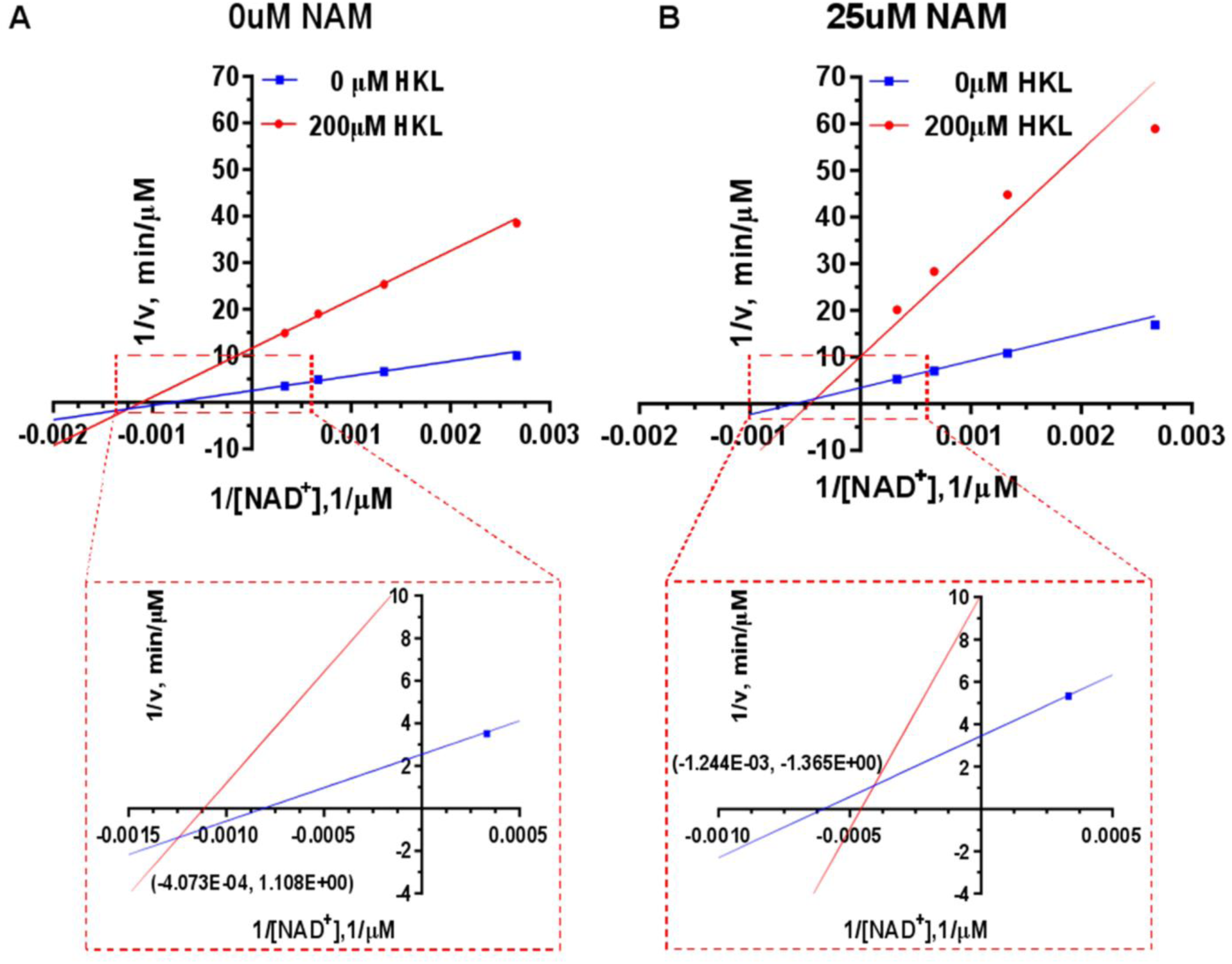
Double reciprocal plots for deacylation initial rate measurements of saturating substrate peptide (FdL2) in the absence and presence of Honokiol with **(A)** 0 µM NAM; **(B)** 25 µM NAM. The zoom in of intersection point with x, y values are provided as insets. [Originally presented in R. Chakrabarti, Mechanism-based enzyme activating compounds, J. Theor. Bio (submitted)]

### False positive testing of labeled assays

**Fig. S7.**
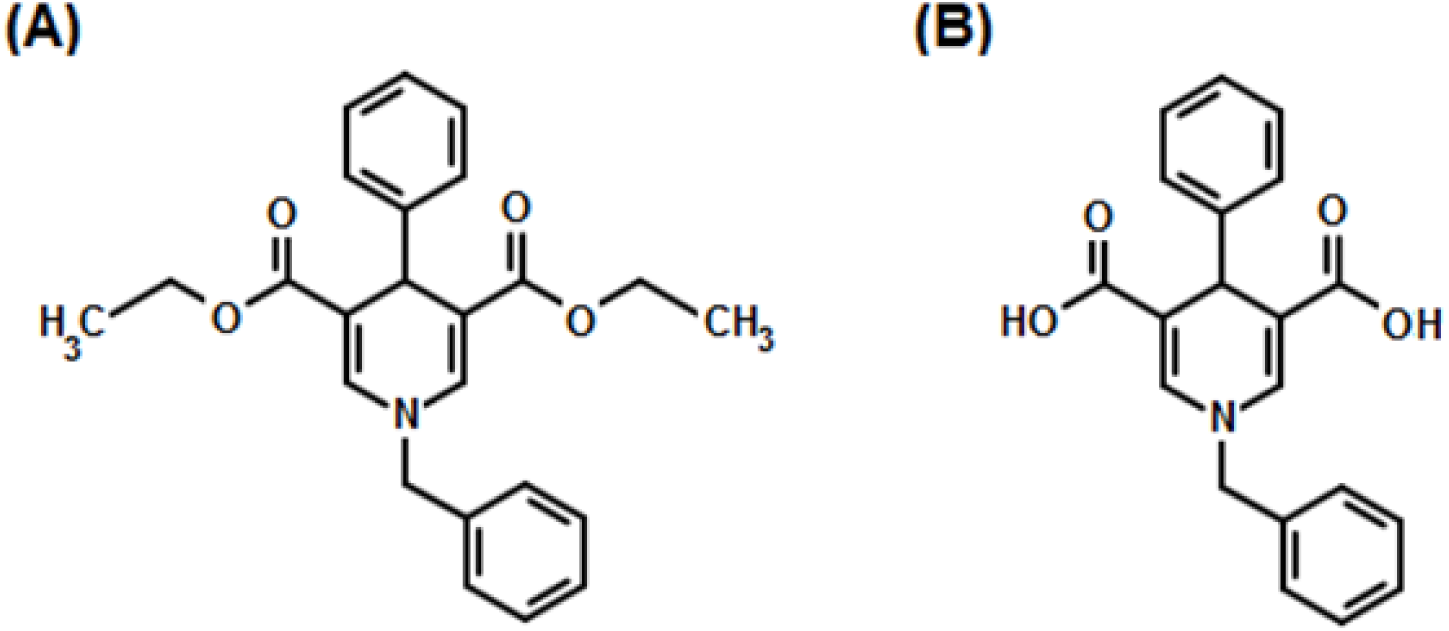
Chemical structures of the sirtuin test compounds. **(A)** N-Benzyl-3, 5-dicar-bethoxy-4-phenyl-1, 4-dihydropyridine (DHP-1). **(B)** N-Benzyl-3, 5-dicarboxy-4-phenyl-1, 4-dihydropyridine (DHP-2).

DHP-1 (Fig. S7(A)) can be dissolved in reaction buffer at 100 µM, but only in the form of a metastable solution. Measurement of DHP-1’s solubility using the above protocol revealed that it is thermodynamically insoluble. By mutating the ester groups in DHP-1 to carboxylic acid groups, we obtain the mutated compound DHP-2 (Fig. S7(B)). In contrast to DHP-1, DHP-2 is thermodynamically soluble (Table B):

**Table B.**
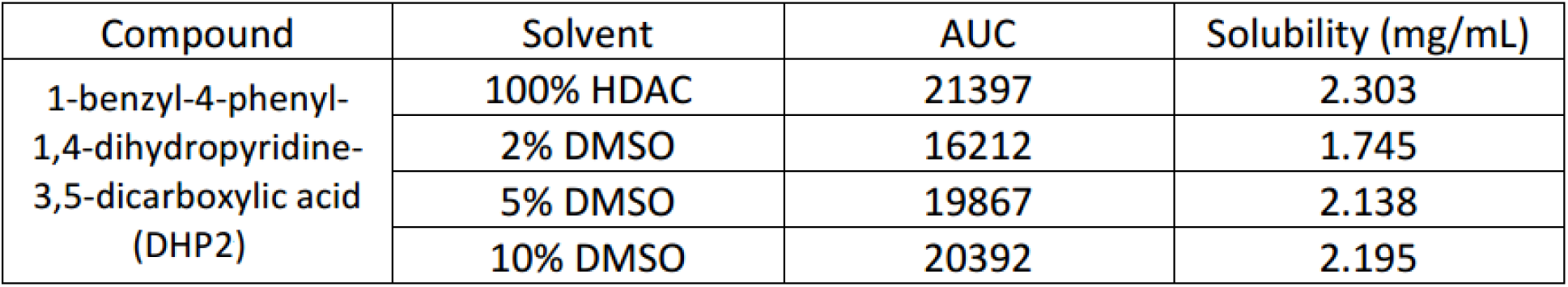
Solubility of DHP2 in different % DMSO/HDAC solution.

The solubility of Honokiol was also assessed using same protocol (Table C):

**Table C.** Solubility of Honokiol in different % DMSO/HDAC solution.

| Compound | Solvent | AUC | Solubility (mg/mL) |
| --- | --- | --- | --- |
| Honokiol | 100% HDAC | n.d. | n.d. |
|  | 1% DMSO | n.d. | n.d. |
|  | 2% DMSO | 260.8 | 0.041 |
|  | 5% DMSO | 766.2 | 0.120 |

Autofluorescence contributes to what is apparently false-positive activation of Sirt3 by the DHP-1 and DHP-2 compounds. A recent study [6, 7] suggested that DHP-1 and 2 were a potent Sirt3 activator using FdL drug discovery kit. However, our results showed that DHP-1 and 2 emit light at wavelengths of 440-460nm, same as the fluorophore (AMC) used in the FdL drug discovery kit (Fig. S8). The strong autofluorescence signal interferes with the readout from the real deacetylation reaction. Standard curves in the presence of different concentrations of DHP-1 (not shown) revealed that (1) the slope changes with varying concentration of [DHP-1]; (2) poor linear correlation was observed at higher [DHP-1]s. Those all indicated that the subtracting baseline fluorescence background did not clear out the autofluorescence effect. Similar effects were observed for DHP-2, where reference [6, 7] reported activation under the conditions tested here (presumably using a single standard curve), whereas we observed apparent inhibition using the FdL assay at these and higher concentrations using a concentration-dependent standard curve. Fig. S9 shows the divergence between dose response curves collected for DHP-2 using a label-free HPLC assay vs the labeled FdL assay. The HPLC assay showed no significant activity modulation at any of the tested concentrations. This divergence was not observed for HKL, which does not display autofluorescence at the detection wavelength of the FdL assay (Fig. S10). Note that the Sirtainty assay used in [7] also displays autofluorescence in the target wavelength range.

**Fig. S8.**
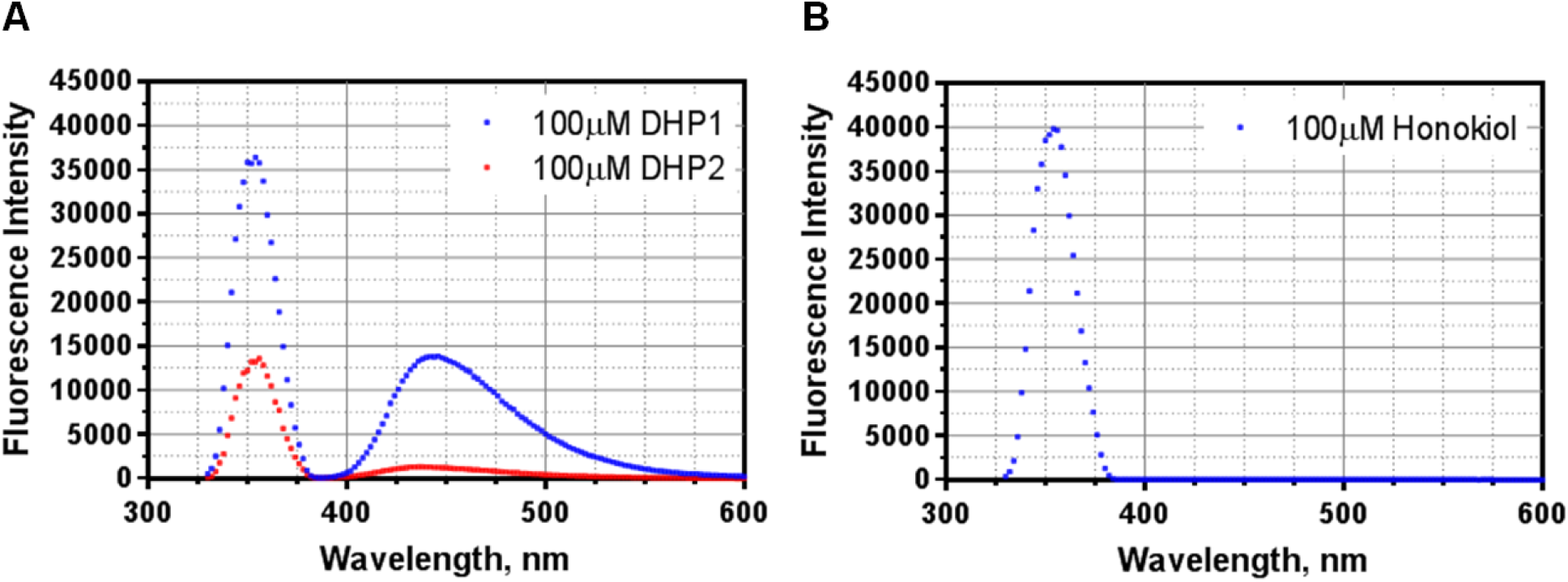
Auto-fluorescence scan for **(A)** DHPs and **(B)** HKL were performed on a multifunctional microplate reader (TECAN Infinite M200 PRO, Switzerland, Tecan Group Ltd.) with setting of Ex=355 nm, Em=330-600 nm, and Gain = 45. The 100 µM of modulator sample solutions were prepared in either DMSO (DHP-1/HKL) or HDAC (DHP-2) based on their solubility (Table **C** and **B**).

**Fig. S9.**
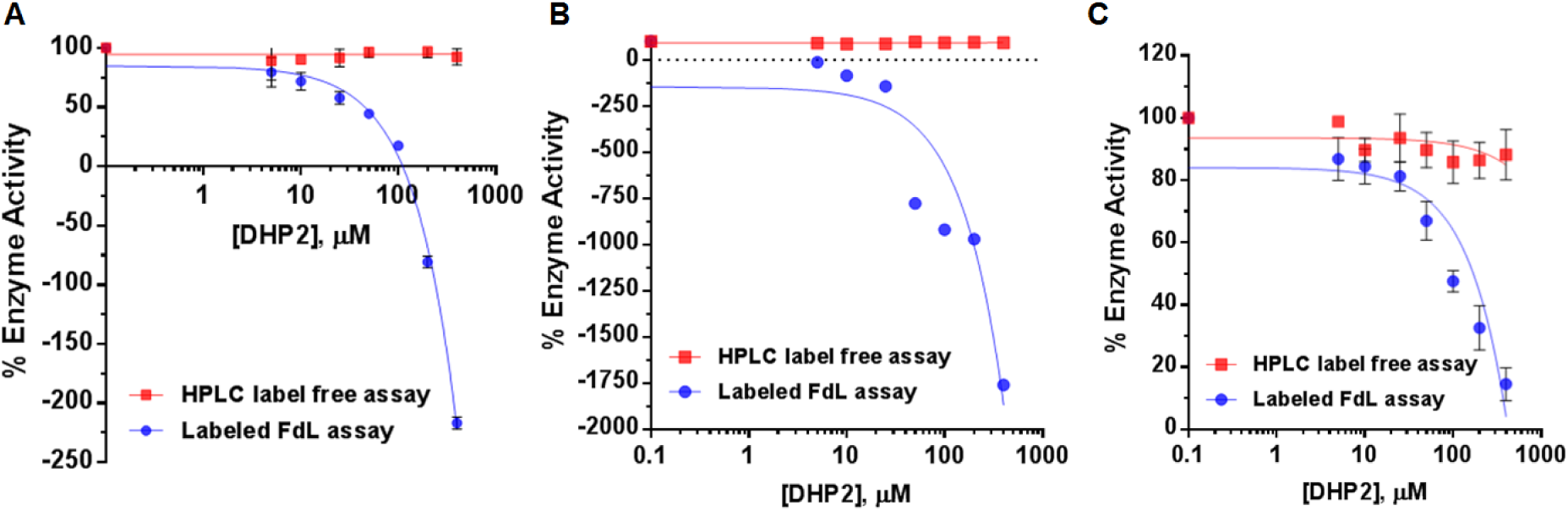
Comparison of dose response of DHP-2 for FdL peptide using label-free HPLC assay and labeled FdL assay. **(A)** Saturating FdL peptide (250µM) and un-saturating NAD^+^ (10µM) (N=2); **(B)** high NAD^+^ (3000µM) and un-saturating FdL peptide (3µM) (N=2); (**C**) 500µM NAD^+^ and saturating FdL peptide (250µM) (N=2) in the presence of different concentrations of DHP-2 range from 0-400 µM using label-free HPLC (red) and labeled FdL assay (blue).

**Fig. S10.**
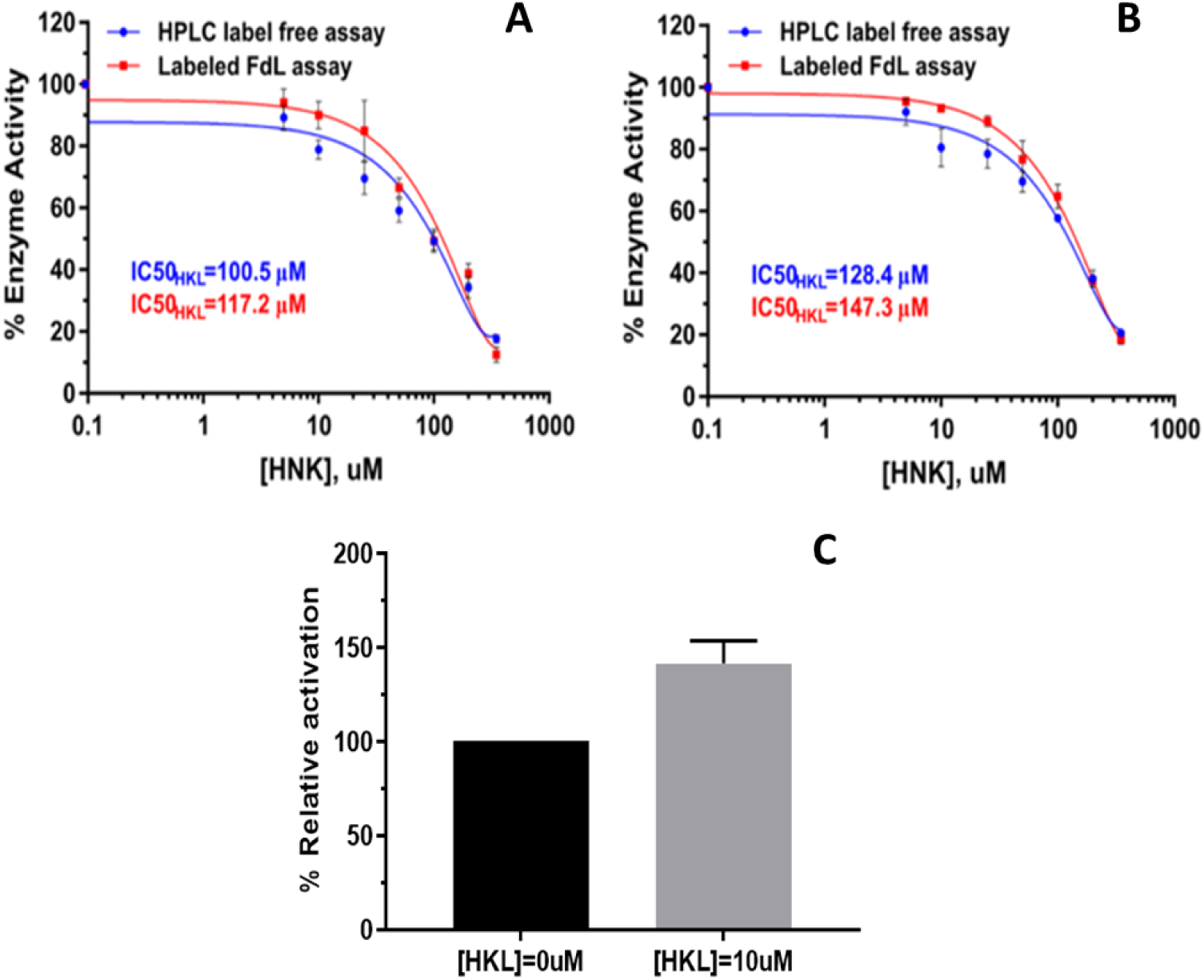
Comparison of dose response of HKL for FdL peptide using label-free HPLC assay and labeled FdL assay. Dose-response curves were measured under conditions where [E]_0_/[S]_0_ << 1, where [S]_0_ denotes the initial concentration of the limiting substrate, in order to maximize the contribution the steady state phase of the reaction to the curve. **(A)** Saturating FdL peptide (250µM) and un-saturating NAD^+^(25µM) (N=2); **(B)** high NAD^+^ (3000µM) and un-saturating FdL peptide (30µM) (N=2) in the presence of different concentrations of HKL range from 0-350 µM using label-free HPLC (red) and labeled FdL assay (blue). **(C)** Activation of SIRT3 in the presence of 10uM HKL at [E]_0_=0.35uM, [NAD^+^]=20uM, [FdL]=10uM, 5 min (N=2).

### Time series fitting

For the MnSOD substrate, as discussed in the text, a single exponential model fit poorly to the complete time series data. The time series data displayed a two phase behavior, one phase between 0 and up to 10 mins. For this system, a single exponential fit would average properties of the first and second phases, which would not provide an accurate steady state rate estimate. Example double exponential fittings are shown in the insets of Fig. 9 and in more detail in Fig. 14.

Also, in order to assess the accuracy of steady state rate calculations for this system, we tested for product inhibition by adding exogenous O-acetylated ADP ribose to the system and evaluating the effect on the time series curve. The results are presented in Fig. S11.

**Fig. S11.**
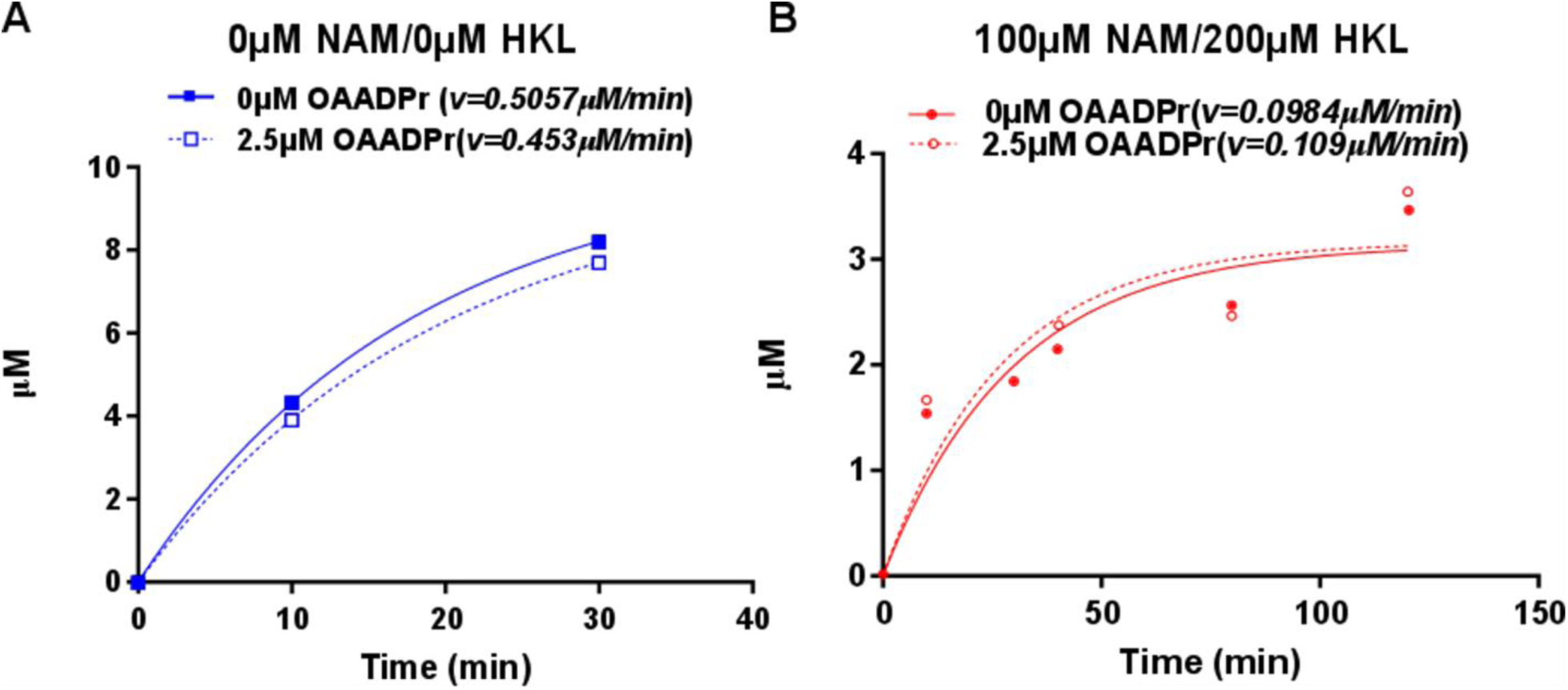
Investigation of OAADPr product inhibition. The deaceylation reactions were performed at [NAD^+^] = 100 µM, [MnSOD K122]= 600 µM in the presence of 2.5 µM OAADPr (A) 0 µM NAM and 0 µM HKL (B) 100 µM NAM and 200 µM HKL. The product formation was monitored by HPLC at desired time points (0, 10, 30, 40, 80, 120 min). The time series were fitted in single exponential function in Prism.

Following system identification of all rate parameters in the steady state model, the rate constant for the limiting chemistry step can be identified based on non-steady state experiments and the analytical solution for the time series dynamics. The time series data and model fittings show that especially at low [NAD^+^], the pre-steady state turnover rates are faster than the turnover rates at steady state and implied by k_cat_/K_m_. This results in the initial deacetylation rates under these conditions being higher in the presence of HKL than in its absence, and significantly higher than the estimated steady-state rates. Note that k_cat_/K_m_ is only applicable under steady state conditions and not at pre-steady conditions such as those at small times or high [E]_0_/[NAD^+^]_0_. The fact that SIRT3/MnSOD has a fast pre-steady state phase means product release rate improvement is a goal of hit to lead evolution.

**Fig. S12.**
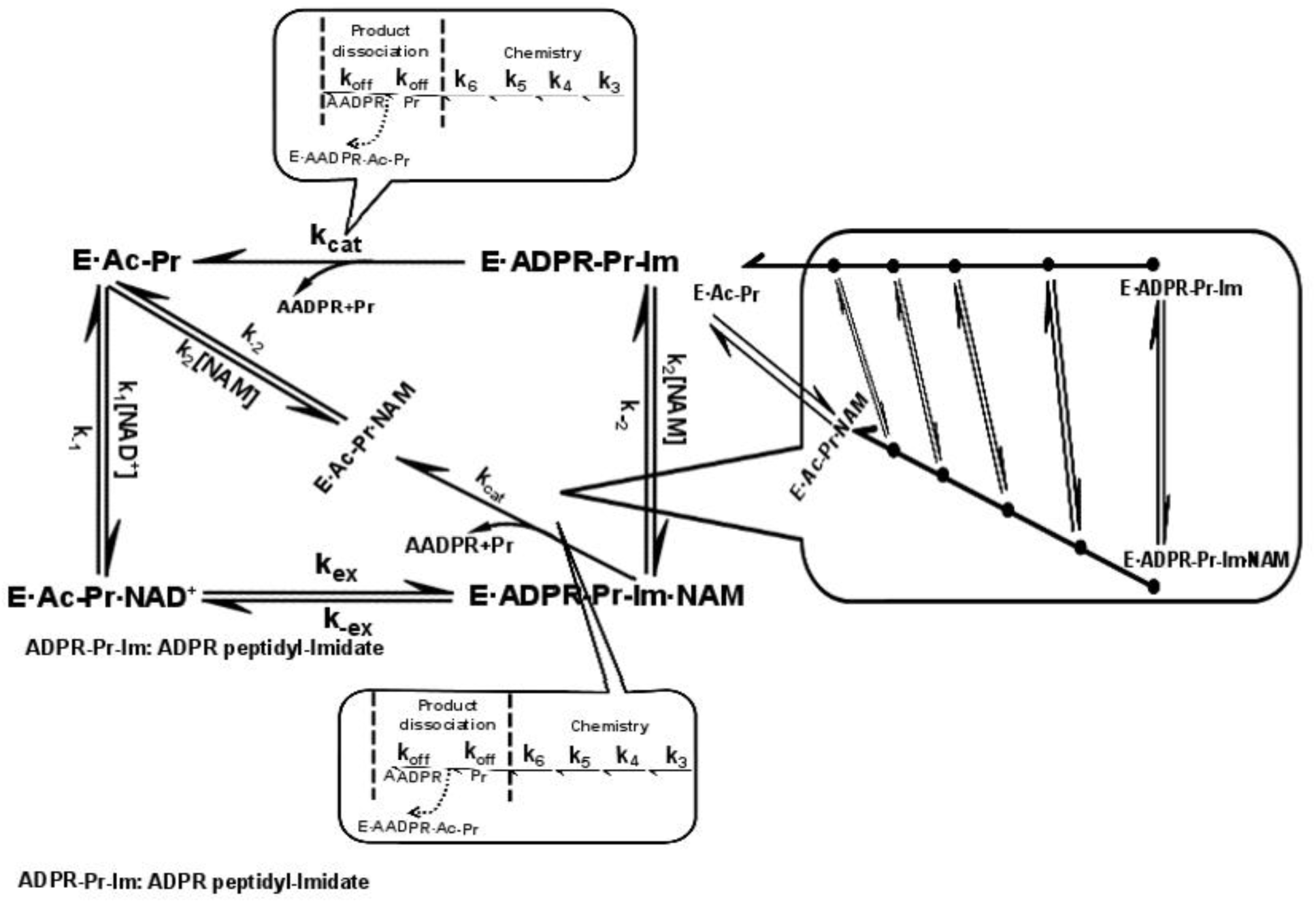
Extended model for sirtuin catalyzed deacetylation including separate representation of elementary deacetylation chemistry and product release steps.

